# Reduced gene templates for supervised analysis of scale-limited CRISPR-Cas9 fitness screens

**DOI:** 10.1101/2022.02.28.482271

**Authors:** Alessandro Vinceti, Umberto Perron, Lucia Trastulla, Francesco Iorio

**Affiliations:** Human Technopole, Viale Rita Levi-Montalcini, 1 - 20157, Milano, Italy; Wellcome Sanger Institute, Wellcome Genome Campus, Hinxton, Cambridge, CB10 1SA, UK

**Keywords:** CRISPR-Cas9 screens, cancer dependency, reference genes, scale-limited screens, supervised analysis

## Abstract

Pooled genome-wide CRISPR-Cas9 screens are furthering our mechanistic understanding of human biology and have allowed us to identify new oncology therapeutic targets. Scale-limited CRISPR-Cas9 screens – typically employing guide RNA libraries targeting subsets of functionally related genes, individual biological pathways, or portions of the druggable genome – constitute an optimal setting for investigating narrow hypotheses and they are easier to execute on complex models, such as organoids and in vivo models. Different supervised methods are used for the computational analysis of genome-wide CRISPR-Cas9 screens; most are not well suited for scale-limited screens as they require large sets of positive/negative control genes (gene templates) to be included among the screened ones. We have developed a computational framework identifying optimal subsets of known essential and nonessential genes (at different subsampling percentages) that can be used as templates for supervised analyses of scale-limited CRISPR-Cas9 screens, while having a reduced impact on the size of the employed library.

**Highlights:**

- Scale-limited CRISPR-Cas9 screens are experimentally easier than genome-wide screens
- Reference gene templates are used for supervised analyses of genome-wide screens
- Reduced templates allow supervised analyses of scale-limited CRISPR-Cas9 screens
- We present optimal reduced templates and a computational method to assemble them

**Graphical abstract:** 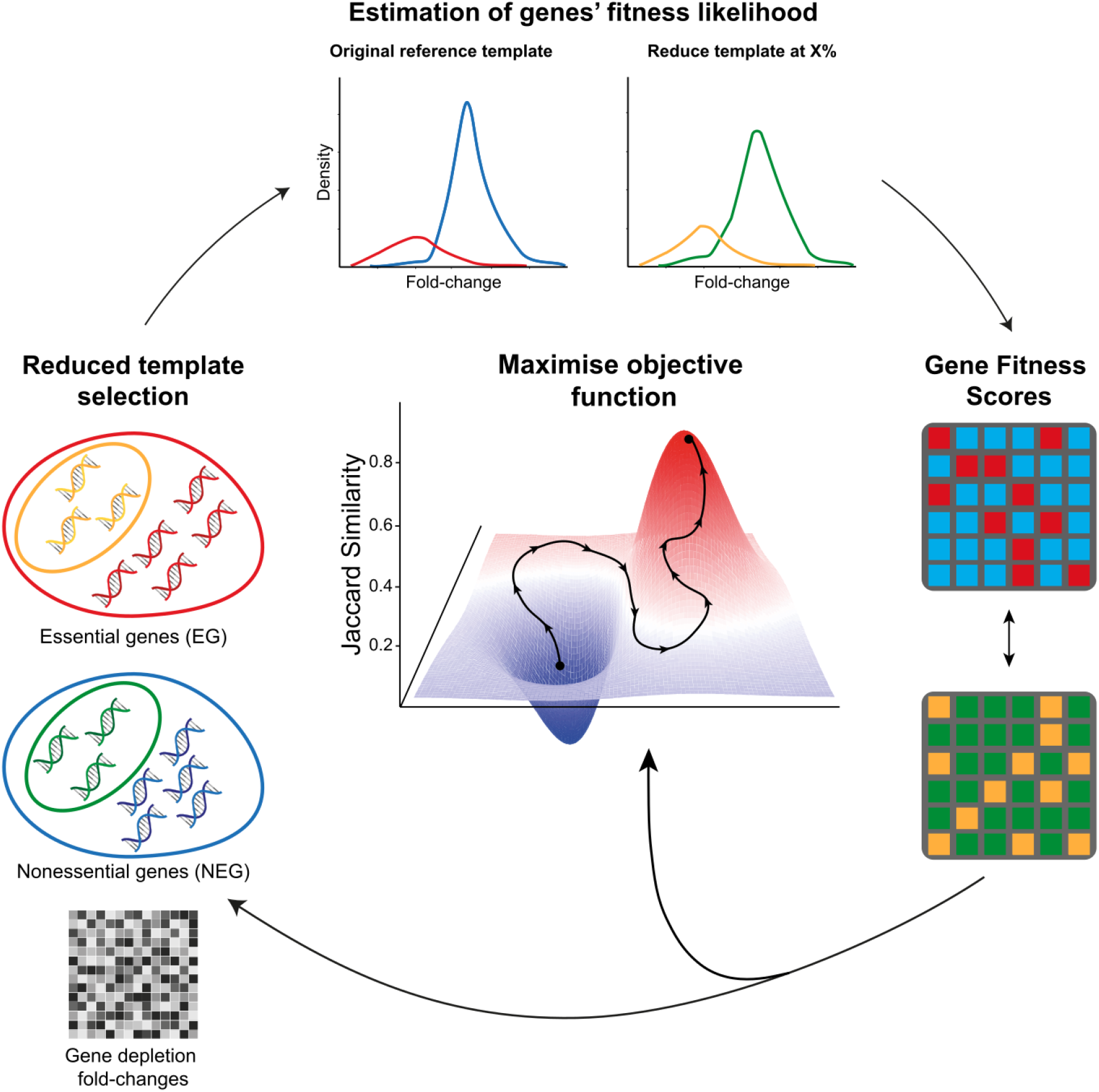

## Introduction

CRISPR-Cas9 knockout (KO) screens are powerful tools that have ushered a new era for genomic research (Shalem et al. 2014; Hartenian and Doench 2015; Barrangou 2020). One of the main successful applications of this technology has been the identification of genes essential for cancer cell survival (i.e. fitness genes) at a genome-scale to support the discovery of new cancer dependencies and therapeutic targets (Behan et al. 2019; Tzelepis et al. 2016; Tsherniak et al. 2017; Meyers et al. 2017), while also widely improving our understanding of human biology (Shalem, Sanjana, and Zhang 2015; Zhou et al. 2014). Focused (or scale-limited) CRISPR-Cas9 knockout screens, employing single-guide RNAs (sgRNAs) libraries of relatively small size, represent a variation of the aforementioned systematic genome-wide approaches in which only a portion of genes are targeted: typically functionally related genes, part of the same pathway or genes coding for druggable proteins (Parrish et al. 2021; Slabicki et al. 2020; Su et al. 2020; Williams et al. 2020; Roesch, OhAinle, and Emerman 2018; Birsoy et al. 2015; Tarumoto et al. 2018; Condon et al. 2021; Girardi et al. 2020; Zhang et al. 2020; Zhu et al. 2021; Wheeler et al. 2020; Turner and Turner 2021).

These scaled-limited screens are commonly used to test narrower biological hypotheses and they require an initial pooled population of cells of significantly smaller size compared to genome-wide screens, in order to obtain appropriate library representation and multiplicity of infection following transfection and selection (Miles, Garippa, and Poirier 2016; Gonçalves et al. 2021; Doench 2018; Peets et al. 2019). For this reason they are also used as an alternative to genome-wide libraries with minimal number of targeting sgRNAs per gene (Gonçalves et al. 2021; DeWeirdt et al. 2020; Peets et al. 2019) to perform mid-scale to quasi genome-wide screens on complex models, such as organoids and in vivo models.

Many analytical tasks performed on data from genome-wide CRISPR-Cas9 screens are either optionally or necessarily performed in a supervised manner. In this scenario, the screening outcomes observed for large sets of positive/negative controls, i.e. genes that are prior known to be essential/nonessential for cell survival, are adopted as benchmark or template classifiers. These outcomes are typically expressed as fold changes (FCs) of targeting sgRNAs’ representation, obtained by contrasting their abundance post-versus pre-applying selective pressure to the screen. These sgRNAs’ FCs are then aggregated at a targeted-gene level yielding *gene depletion FCs*.

The analytical tasks accomplished by supervised methods range from quality control, to FC scaling for inter-screen comparisons and interpretability, to calling statistical significant essential genes. For the latter task, several tools have been proposed (Li et al. 2014; Traver Hart and Moffat 2016; Kim and Hart 2021; Doench et al. 2016) with the most widely applied ones being the unsupervised method MAGeCK (Model-based Analysis of Genome-wide CRISPR-Cas9 Knockout) (Li et al. 2014, 2015) and the supervised method BAGEL (Bayesian Analysis of Gene EssentiaLity) (Kim and Hart 2021; Traver Hart and Moffat 2016), and BAGEL outperforming MAGeCK when applied to data from CRISPR-induced phenotypes of weak to middle intensities (Behan et al. 2019).

In particular, BAGEL (Traver Hart and Moffat 2016; Kim and Hart 2021) uses two sets of, respectively, prior known essential and nonessential genes (EGs and NEGs respectively) as a template classifier (**Fig. 1A**). First, it estimates two probability density functions (pdfs) from the depletion FCs of the sgRNAs targeting the EGs and the NEGs, respectively. Then, for each sgRNA targeting any other gene, it computes a Bayes factor (BF) quantifying the likelihood of the observed sgRNA’s FC being generated by the EG pdf over that of it being generated by the NEG pdf. Gene-level BFs are then computed by aggregating individual sgRNA-level scores on a targeted gene basis, for example by summing them up or taking median or mean values.

**Fig. 1.**
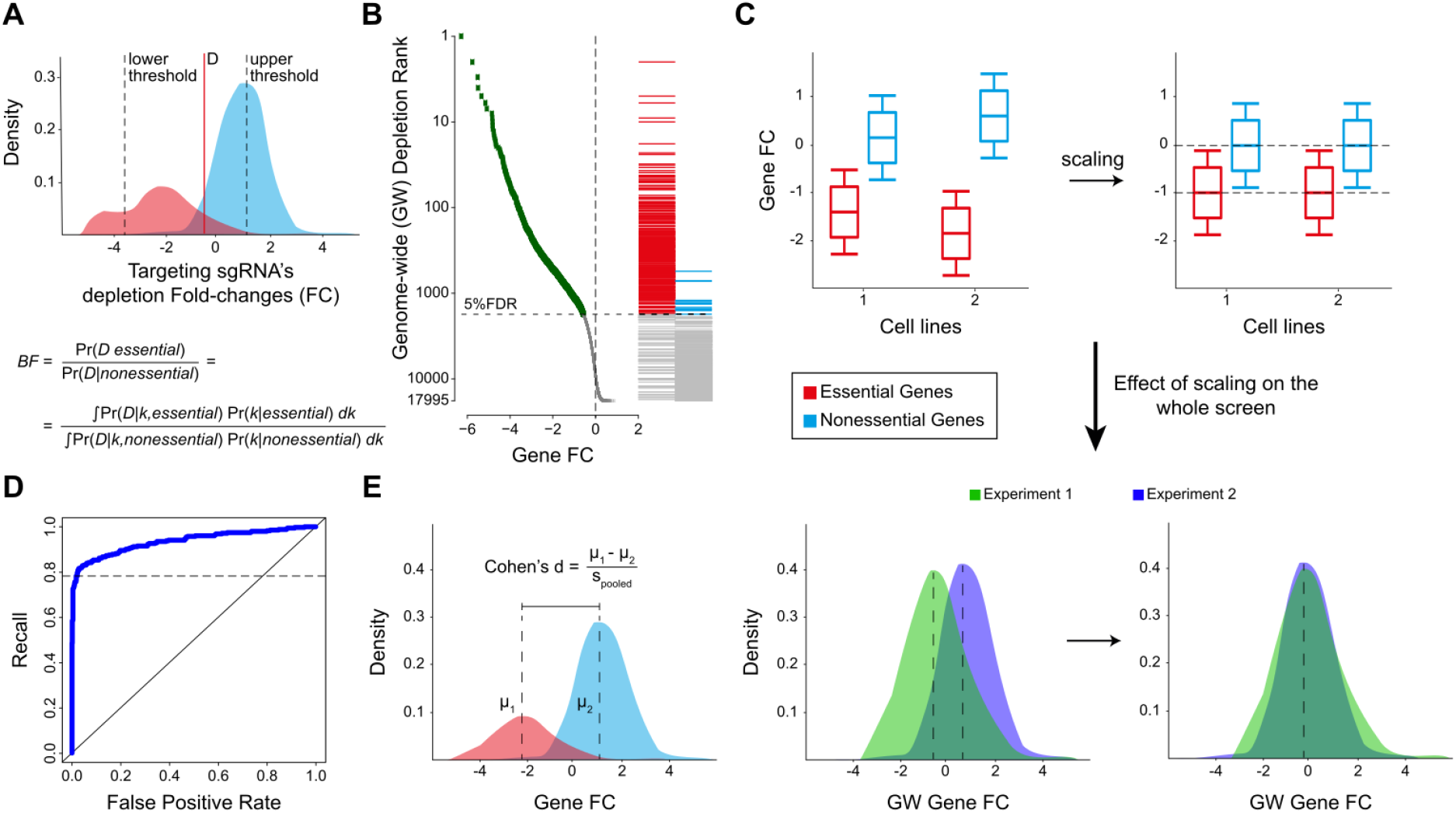
Supervised analysis of CRISPR-Cas9 knockout screens using reference gene templates. **A**. Bayesian Analysis of Gene EssentiaLity (BAGEL) method for the estimation of gene essentiality significance. BAGEL estimates two probability density functions (pdfs) from the depletion fold changes (FCs) of the single-guide RNAs (sgRNAs) targeting prior known essential and the nonessential genes (EGs and NEGs), respectively. Then, for each sgRNA targeting any other gene, it computes a Bayes factor (BF) quantifying the likelihood of the observed sgRNA’s depletion FC being generated by the EG pdf over that of it being generated by the NEG one. **B**. False Discovery Rate (FDR) method for the estimation of gene essentiality significance. The gene-level FCs are ranked in increasing order, then FDRs of positive/negative essential genes used as a template are computed when considering as predicted essentials all the genes from the beginning of the ranked list to each possible rank position, one FDR value per rank position. The genes above the rank position corresponding to an FDR less or equal to a given percentage (typically 5% or 10%) are called significantly essential at that percentage of FDR. **C**. scaling of gene depletion FCs derived from two screens. After the scaling, the FC distribution of sgRNAs targeting the EGs will have a median of -1, and that for the NEGs will have a median of 0 (upper plots), and the gene essentiality distributions from different screens will be aligned, comparable and more interpretable (lower plots). **D**. Quality control assessment of a screen: a profile of gene depletion FCs is used as a rank classifier of positive/negative controls, i.e. the EGs and NEGs respectively. Then the screen quality is assessed by means of Receiver Operating Characteristic analysis. **E**. An alternative metric of screen quality using a gene template is the difference in mean/median depletion FCs between the positive and negative control sets with respect to their pooled standard deviation, expressed as Cohen’s d coefficient. The higher the coefficients, the higher the separation of the EGs/NEGs’ FCs, thus the screen quality.

Another method to call significantly essential genes, based on a required false discovery rate (FDR) level (**Fig. 1B**), consists in ranking all the gene-level FCs in increasing order (or BAGEL BFs in decreasing order), then computing FDRs of positive/negative essential gene controls, considering as predicted essentials all the genes from the beginning of the ranked list to each possible rank position, one FDR value per rank position. Genes above the rank position corresponding to an FDR less or equal to a given percentage (typically 5% or 10%) are then called significantly essential at that percentage of FDR (Behan et al. 2019; Pacini et al. 2021; Dempster, Pacini, et al. 2019; T. Hart et al. 2015).

Reference gene templates are also used for other purposes. For instance, they are employed to scale gene essentiality profiles such that the median depletion FC of EGs and NEGs will be equal to -1 and 0, respectively (**Fig. 1C**) (Aguirre et al. 2016; Munoz et al. 2016; Pacini et al. 2021; Dempster, Pacini, et al. 2019; Meyers et al. 2017; Tsherniak et al. 2017). This allows gene essentiality profiles to be more interpretable and comparable when derived from different screens, or from screening different models. Finally, reference gene templates of EGs and NEGs are routinely used as positive/negative controls to evaluate penetrance of the CRISPR-Cas9 phenotype and screening data quality. This is usually quantified via Receiver Operating Characteristic (ROC) analysis, assessing the performances of a genome-wide profile of gene depletion FCs when considered as rank based classifiers of the control gene-sets (**Fig. 1D**) (Traver Hart et al. 2014; T. Hart et al. 2015; Tzelepis et al. 2016; Behan et al. 2019; Gonçalves et al. 2021; Koike-Yusa et al. 2014). As an alternative, the difference in mean depletion FCs for the positive/negative control sets with respect to their pooled standard deviation (computed via the Cohen’s d or Glass Deltas) (**Fig. 1E**) is also used to score the CRISPR-Cas9 induced phenotype intensity (Behan et al. 2019; Kim and Hart 2021).

The smallest gene template available to date of reference EGs and NEGs is from a recent release of BAGEL (Traver Hart et al. 2014; Steinhart et al. 2017) and consists of 1,611 genes (684 EGs and 927 NEGs, **Table 1**). Hence, to use BAGEL or other aforementioned supervised tools for the analysis of a CRISPR-Cas9 screen these genes need to be forcibly included among those targeted by the employed library. This has a significant impact on the scale of the experiment to perform and it proves quite prohibitive for scale-limiteds screens, where the number of reference genes would become comparable or even larger than that of the genes under investigation.

**Table 1.** State-of-the-art sets of reference essential and nonessential genes widely used by the community and considered in this study to derive templates of reduced size.

| Set name | Set Type | Description and Source | Dataset of origin and derivation method |
| --- | --- | --- | --- |
| *Hart2014* | Set of nonessential genes | A set of 927 genes presented in (Traver Hart et al. 2014). | Large collection of shRNA gene dependency profiles analysed with a linear algebra approach. |
| *Hart2017* | Set of essential genes | A set of 684 genes introduced in (Traver Hart et al. 2017). | BAGEL reanalysis of 17 genome-scale knockout screens in human cell lines performed with different libraries. |
| *ADaM2021* | Set of essential genes | A set of 1075 genes presented in (Vinceti et al. 2021). | ADaM analysis of a large collection of gene dependency profiles of CRISPR-Cas9 screens of 855 human cancer cell lines from different tissue-lineages/cancer-types, integrating Project Score and Project Achilles 20Q2 (Dwane et al. 2021; Dempster, Pacini, et al. 2019; Pacini et al. 2021). |

Here, we present a computational framework for assembling gene templates of reduced size allowing supervised analyses of data from scale-limited CRISPR-Cas9 screens, while having a limited, user-defined impact on the overall library size. Our method, the *Minimal Template Estimator* (MinTEs), is based on a greedy optimisation algorithm. More specifically, using large public available cancer dependency map (DepMap) datasets assembled via genome-wide CRISPR-Cas9 screens of immortalised human cancer cell lines, MinTEs optimises an objective function to identify sgRNAs targeting increasingly smaller informative subsets of state-of-the-art reference EGs and NEGs (from now, the reference gene-sets). MinTEs is trained on two large genome-wide pooled CRISPR-Cas9 datasets from Project Score (Dwane et al. 2021) and project Achilles (Meyers et al. 2017), respectively. These datasets have been obtained using popular genome-wide sgRNA libraries: the Sanger’s Kosuke Yuka (KY) library (Tzelepis et al. 2016) for Project Score, and the AVANA library (Doench et al. 2016) for project Achilles.

When used in supervised analyses of CRISPR-Cas9 screens, the resulting library-specific reduced templates (RTs) produce BF scores (thus sets of predicted fitness genes) that are the closest possible to those yielded when using whole state-of-the art reference gene-sets. We have used MinTEs to generate KY- and AVANA-specific RTs, as well as library-independent RTs. We have tested the performances of library-specific RTs, as well as library-independent RTs on data generated with independent libraries, and observed improved performances with respect to random subsampling. In addition, we have observed strikingly better performances also with respect to a data driven subsampling strategy, where the reduced templates are assembled picking genes that are never essential or always essential (respectively for NEGs and EGs control sets) in large DepMap datasets. Finally, we also benchmarked the reliability of the RTs when employed to estimate gene essentiality with the FDR method and in other supervised use-case analyses, showing large outcome concordance with respect to using the original reference gene-sets.

## Results

### Overview of the Minimal Template Estimator framework

Minimal Template Estimator (MinTEs) is a a greedy search computational framework for the identification of optimal subsets of essential (EGs) and nonessential genes (NEGs) to be used in supervised analysis of scale-limited CRISPR-Cas9 screens (**Fig. 2A**). These subsets are derived from state-of-the-art reference sets (STAR *Methods*), at different percentages of subsampling, from 95% down to 5% of the reference gene-sets’ size (**Table 2**). This yields a flexible portfolio of gene pools, with sizes ranging from hundreds to few tens, which users can select based on the desired library size while setting up a new scale-limited screen.

**Table 2.** Size of the reduced gene templates with respect to the percentage of subsampling and considered reference set (i.e. ADaM2021 or Hart2017).

| Percentage | n. EG (ADaM2021) | n. EG (Hart2017) | n. NEG |
| --- | --- | --- | --- |
| 5% | 54 | 34 | 46 |
| 10% | 108 | 68 | 93 |
| 15% | 161 | 103 | 139 |
| 20% | 215 | 137 | 185 |
| 25% | 269 | 171 | 232 |
| **30%** | 322 | 205 | 278 |
| **35%** | 376 | 239 | 324 |
| **40%** | 430 | 274 | 370 |
| **45%** | 484 | 308 | 417 |
| **50%** | 538 | 342 | 463 |
| **55%** | 591 | 376 | 509 |
| **60%** | 645 | 410 | 556 |
| **65%** | 699 | 445 | 602 |
| **70%** | 753 | 479 | 648 |
| **75%** | 806 | 513 | 695 |
| **80%** | 860 | 547 | 741 |
| **85%** | 914 | 581 | 787 |
| **90%** | 968 | 616 | 833 |
| **95%** | 1021 | 650 | 880 |
| **100%** | 1075 | 684 | 926 |

**Fig. 2.**
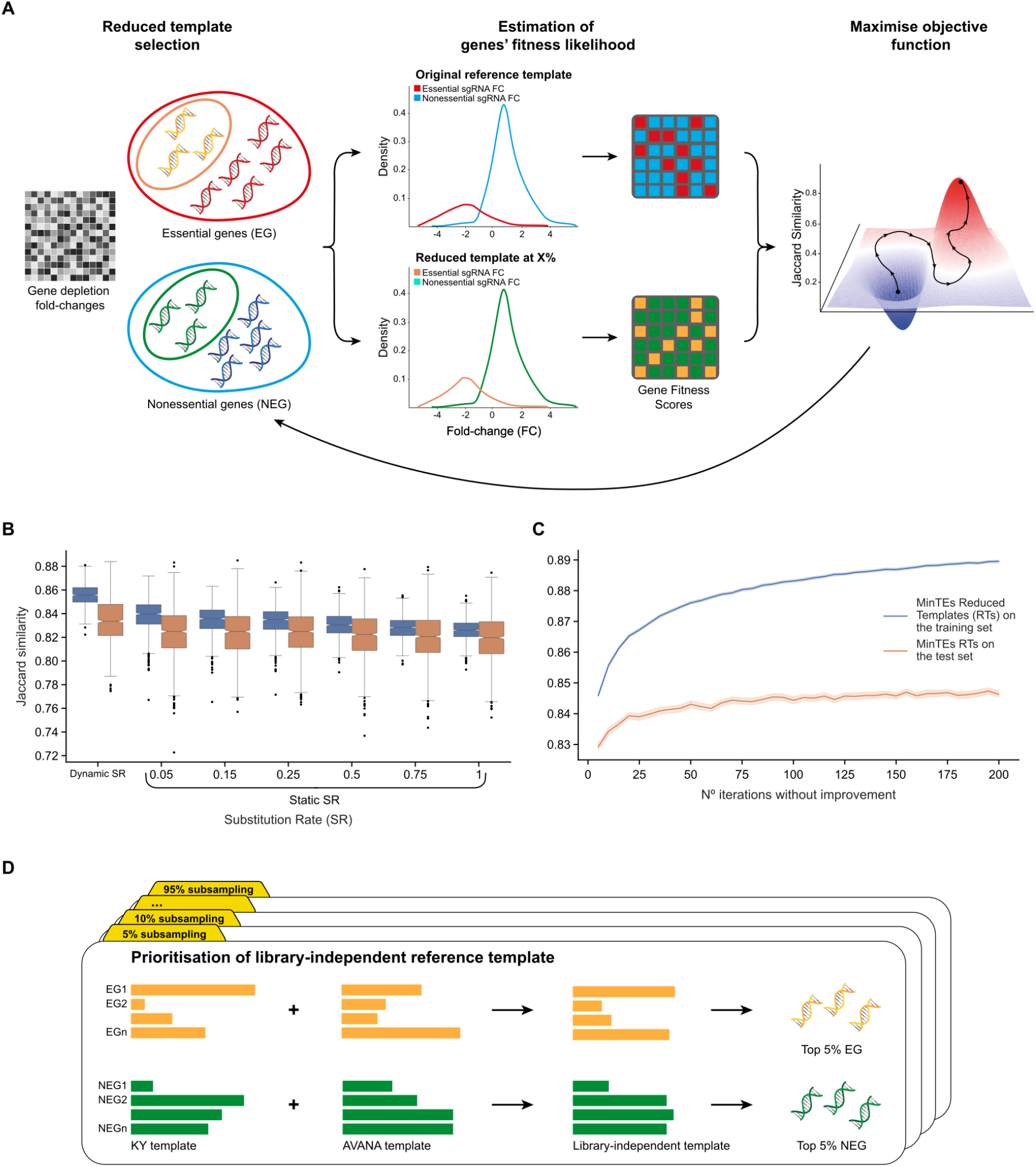
Overview of the MinTEs computational framework. **A**. MinTEs’s goal is to identify subsets of reference essential/non-essential genes, i.e. a reduced gene templates (RT), such that, when used for the analysis of CRISPR-Cas9 screens with a supervised method, this RT yield sets of predicted fitness genes that are the closest as possible to those obtained when using the original reference gene-sets. This is achieved by optimising an objective function defined as the average Jaccard similarity score between fitness gene-sets predicted with whole template or a candidate RT when applying the supervised method BAGEL to a large collection of public gene dependency profiles. This optimisation is executed across 1,000 train-test sets reshufflings and the best RTs are selected based on a prioritisation strategy considering their performance across reshuffling trials. This produces RTs that are specific to the CRISPR-Cas9 single guide RNA library used to generate the gene dependency profiles used in input. **BC**. Average Jaccard similarity scores between fitness gene-sets predicted by BAGEL when using as classification templates the MinTEs derived RTs and the Hart2017 essential and Hart2014 nonessential gene-sets on the Project Score dataset (obtained with the KY library) at 5% percentage of subsampling and testing different instances of substitution rate (SR) (x-axis) (**B**) and *C* (i.e., n° iterations without improvement, *STAR* method) (x-axis) (**C**). RTs’ performances are split between training and test sets (different colours). We compared static vs dynamic SRs, the latter following an exponential decay function (*STAR* method). **D**. To assemble library-independent RTs, each library-specific RT is tested on a CRISPR-Cas9 dataset generated with an independent library. Then, a library-independent score is computed by aggregating and weighting the performances of each RTs across CRISPR-Cas9 datasets using an ad-hoc prioritisation strategy. The top-performing RT is then selected and put forward as the library-independent one.

MinTEs starts from the analysis of public CRISPR-Cas9 datasets derived from screening hundreds of immortalised human cancer cell lines, with genome-wide sgRNA libraries (Dwane et al. 2021; Dempster, Rossen, et al. 2019; McFarland et al. 2018) (STAR *Methods*). Here we present library-specific results (**Table S1**,**S2**) - obtained using in input the Project Score dataset (Dwane et al. 2021; Behan et al. 2019) (obtained with the the Sanger’s Kosuke Yuka (KY) library (Tzelepis et al. 2016)), and the project Achilles dataset (Dempster, Pacini, et al. 2019) (obtained with the AVANA library (Doench et al. 2016)) - as well as library-independent results (**Table S3**) assembled through a prioritisation strategy based on the performances of the library-specific RTs.

In MinTEs the CRISPR-Cas9 dataset in input is first split into a training set and a test set, then random subsets of EGs and NEGs, whose size depends on the desired percentage of subsampling, are initialised. These randomly selected reduced templates (RTs) are used in BAGEL (Kim and Hart 2021) for the estimation of fitness genes across the screened cell lines in the training set. The predicted fitness genes are then compared to those obtained when running BAGEL with the whole reference gene-sets. This yields an average Jaccard similarity score: the higher this score, the larger the agreement between the predictions obtained with full reference sets and those obtained with the RT under consideration.

An iterative RT optimisation strategy is then followed by MinTEs, where the Jaccard score of a new RT candidate (obtained from the previous one by randomly replacing individual genes) is compared to that obtained in the previous step, and the RT with higher Jaccard score is iteratively retained until a plateau is reached. Across iterations, a cooling schedule (i.e. a function defining a finite number of transition states to guide the algorithm to convergence) increasingly lowers the number of gene swapping between a candidate RT and that obtained in the previous optimisation step (**Fig. 2BC** and STAR *Methods*). Upon convergence of the Jaccard scores, the performances of the optimised RT are assessed on the test set. This is executed across 1,000 train-test reshufflings for each combination of parameters (i.e. dataset in input, starting reference EG set and percentage of subsampling) and it is followed by a prioritisation strategy selecting the best RT across train-test reshufflings (STAR *Methods*).

Initially, we explored different settings for our computational framework (**Fig. S1**). First, we tested different heuristic optimisation strategies (**Fig. S1AB**), namely simulated annealing and a genetic algorithm with fixed mutation and crossover rates. Although these methods performed generally better in the training phase, they did not offer substantial improvements on the test set (**Fig. S1A**), while requiring a larger number of iterations to achieve convergence (**Fig. S1B**). Therefore, we decided to utilise and pipeline the greedy search strategy also in the light of possible periodic recomputations of the RTs as larger (or derived from more complex models) training sets will become increasingly available. Within this framework, the use of BAGEL for assessing the quality of RTs selection offers a major advantage: it is the most elaborated supervised analytical method among those we listed, and it allows to clearly define an objective function to maximise while assembling the RTs. BAGEL also shows greater sensitivity to gene-sets subsampling, making it suitable to be employed in a controlled stepwise improvement strategy. Secondly, we tested other popular similarity metrics as alternatives to the Jaccard similarity (**Fig. S1C**, specifically true positive rate, positive predicted value and F1 score). We obtained similar results with respect to the Jaccard similarity, with MinTEs selected RTs systematically outperforming random RTs.

We applied MinTEs to compute library-specific RTs at different percentages of subsampling of the original reference gene-sets, for the AVANA and KY libraries (**Fig. 2D**, and **Table S1**,**S2**), and using two different reference EG sets (**Table 1**): the Hart2017 set - used by the latest version of BAGEL (Traver Hart et al. 2017) - and the ADaM2021 set, which we have shown to better recapitulate oncogenetic and other context-specific dependencies than the Hart2017, when used for the analysis of large cancer dependency datasets (Vinceti et al. 2021).

Finally, we assembled library-independent RTs. First, we cross-tested library-specific RTs (obtained from a given reference set of EGs and a fixed percentage of subsampling) on the never-before-seen CRISPR-Cas9 datasets (e.g. testing KY-specific RTs the Achilles datasets - obtained with the AVANA library - and vice versa). Then we applied a prioritisation strategy (STAR *Methods*) to select the top-performing library-independent RTs across cross-tests (**Table S3**).

### Performance assessment of library-specific reduced templates

We assembled library-specific reduced templates (RTs) for the Sanger KY (Tzelepis et al. 2016) and AVANA (Doench et al. 2016) libraries, applying MinTEs to cancer DepMap data from project Score (Dwane et al. 2021) and project Achilles - release 21Q2 (Dempster, Rossen, et al. 2019) - respectively, and using two different starting reference sets of essential genes (EGs) - the Hart2017 (Traver Hart et al. 2017) and ADaM2021 (Vinceti et al. 2021) - across subsampling percentages ranging from 95% to 5%, with a 5% step decrease. We computed the performances of each RT in terms of similarity of fitness gene-sets predicted by BAGEL when using as a template the RT under consideration and the reference EGs/NEGs sets from which it was derived, respectively. This was quantified using a Jaccard similarity score, averaged across all screened cell lines in the input cancer DepMap dataset. For comparison, we also computed performances of randomly subsampled RTs (1,000 samplings per tested subsampling percentage, **Fig. S2**) and DepMap-driven RTs. The random RTs were obtained by selecting random subsets of reference gene-sets, with every gene having the same probability to be selected, according to the subsampling percentage under consideration. On the other hand, the DepMap-driven RTs were obtained by selecting, from the cancer DepMap datasets, a subset of genes with the lowest and highest average FCs, as EGs and NEGs respectively, per each subsampling percentage.

Overall, the library-specific MinTEs RTs outperformed the random RTs and the DepMap-driven RTs, consistently across libraries, starting reference gene-sets, and subsampling percentages (**Fig. 3, S2** and **S3**, STAR *Methods*). Indeed, the gap in performance was strikingly higher with respect to random RTs, when considering smaller percentages of subsampling of the reference gene-sets. In particular, the 5% RTs yield Jaccard similarity scores greater than 0.85 in all cases but one (i.e. RTs derived from Hart2017 as reference essential gene-set and tested on the dataset generated with the AVANA library), demonstrating the ability of the smallest RTs to retain the vast majority of fitness genes identified by the whole original templates. This suggests that the RTs would be particularly beneficial for scale-limited CRISPR-Cas9 screens.

**Fig. 3.**
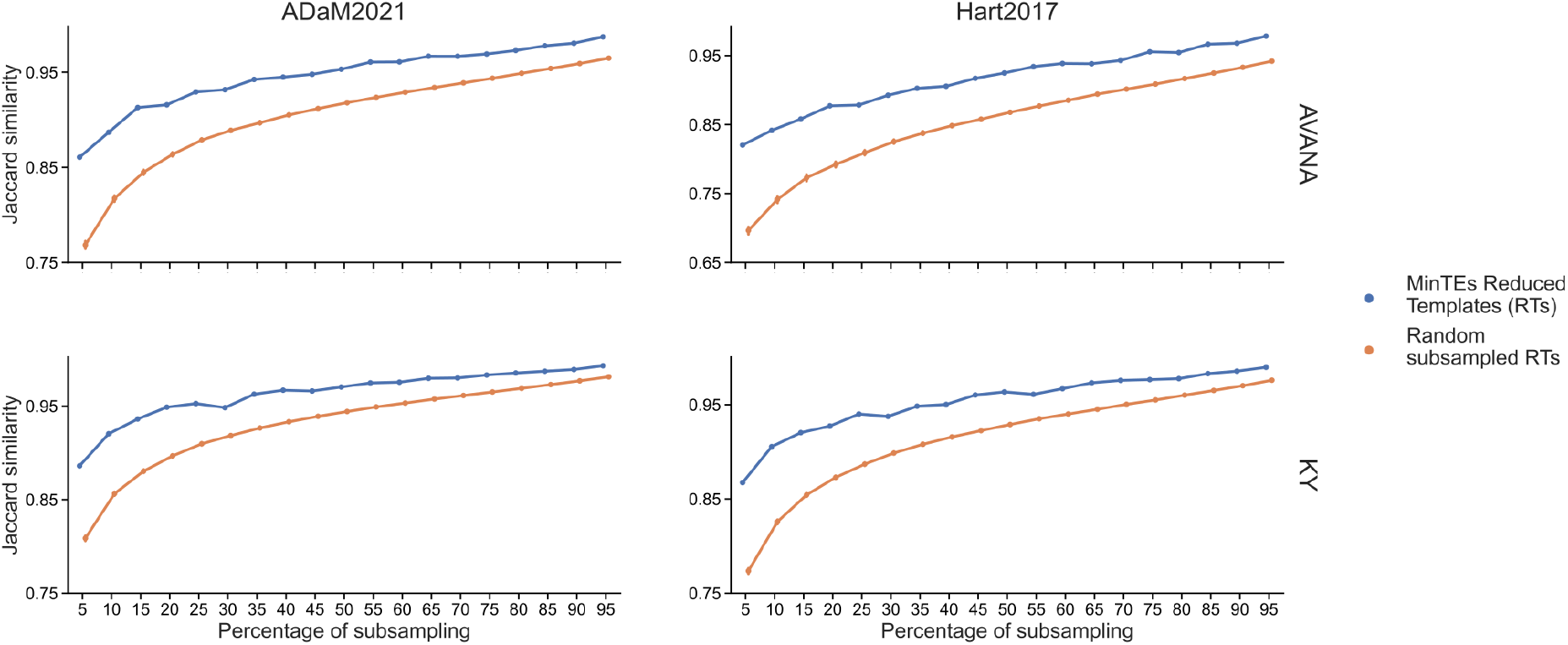
Performances of MinTEs library-specific reduced templates (RTs). Average Jaccard Similarity scores between fitness gene-sets predicted by BAGEL when using as classification templates the MinTEs derived RTs or randomly selected RTs (as indicated by the different colours) obtained from two state-of-the-art reference essential gene-sets (different columns) and the same reference set of nonessential genes, for two different single guide RNA libraries (different rows), across percentage of subsampling. For random performances, the bars show 95% confidence interval computed across 1,000 train-test reshufflings of the input dataset for each combination of parameters (i.e. reference gene-set, CRISPR-Cas9 library and percentage of subsampling).

### Validation of library-independent reduced templates using independent datasets

We tested the performances of the MinTEs derived library-independent reduced templates (RTs) on both cancer dependency datasets used to derive them, as well as on three additional datasets derived by CRISPR-Cas9 screening the HT-29 cell line with independent libraries: respectively, Brunello (Sanson et al. 2018), GeCKOv2 (Sanson et al. 2018), and MiniLibCas9 (Gonçalves et al. 2021). We performed our assessment showcasing typical supervised analyses summarised in **Fig. 1**.

#### Use case A: Estimation of gene essentiality using BAGEL

For every combination of initial reference gene-set (ADaM2021 or Hart2017), tested library (AVANA, KY, Brunello, GeCKOv2 or MiniLibCas9) and percentage of subsampling (95% to 5%), we computed average (for the datasets generated with AVANA and KY, encompassing multiple cell line screens) or individual (for the HT-29 screen, obtained with the independent libraries) Jaccard scores as detailed in the previous section, i.e. quantifying the similarity between BAGEL outputted fitness gene-sets when used with whole reference gene-sets versus RT.

We observed higher Jaccard scores between whole reference gene-sets and MinTEs RT predictions compared to those between whole reference gene-sets and randomly selected RT predictions, for every combination of tested parameters both on the training and validation libraries (**Fig. S4**), despite profound differences in the tested libraries (such as in the average numbers of targeting sgRNAs per gene and targeted sequence positions), as well as experimental conditions across compared screens (such as in screening time lengths).

Particularly, for the independent libraries, Jaccard scores were above 0.8 for the vast majority of tested subsampling percentages (**Fig. S4**), thus indicating that the MinTEs library-independent RTs are able to retain a faithful representation of predicted fitness genes with respect to the original reference gene-sets, when used as classification template with BAGEL. The advantage of using library-independent RT sets can be especially appreciated when focusing on minimal percentages of subsampling, such as 5%, providing a substantial reduction in gene-set size and much higher performances compared to random subsampling (**Fig. 4A**,**B**), as well as DepMap-driven RTs (**Fig. S5**).

**Fig. 4.**
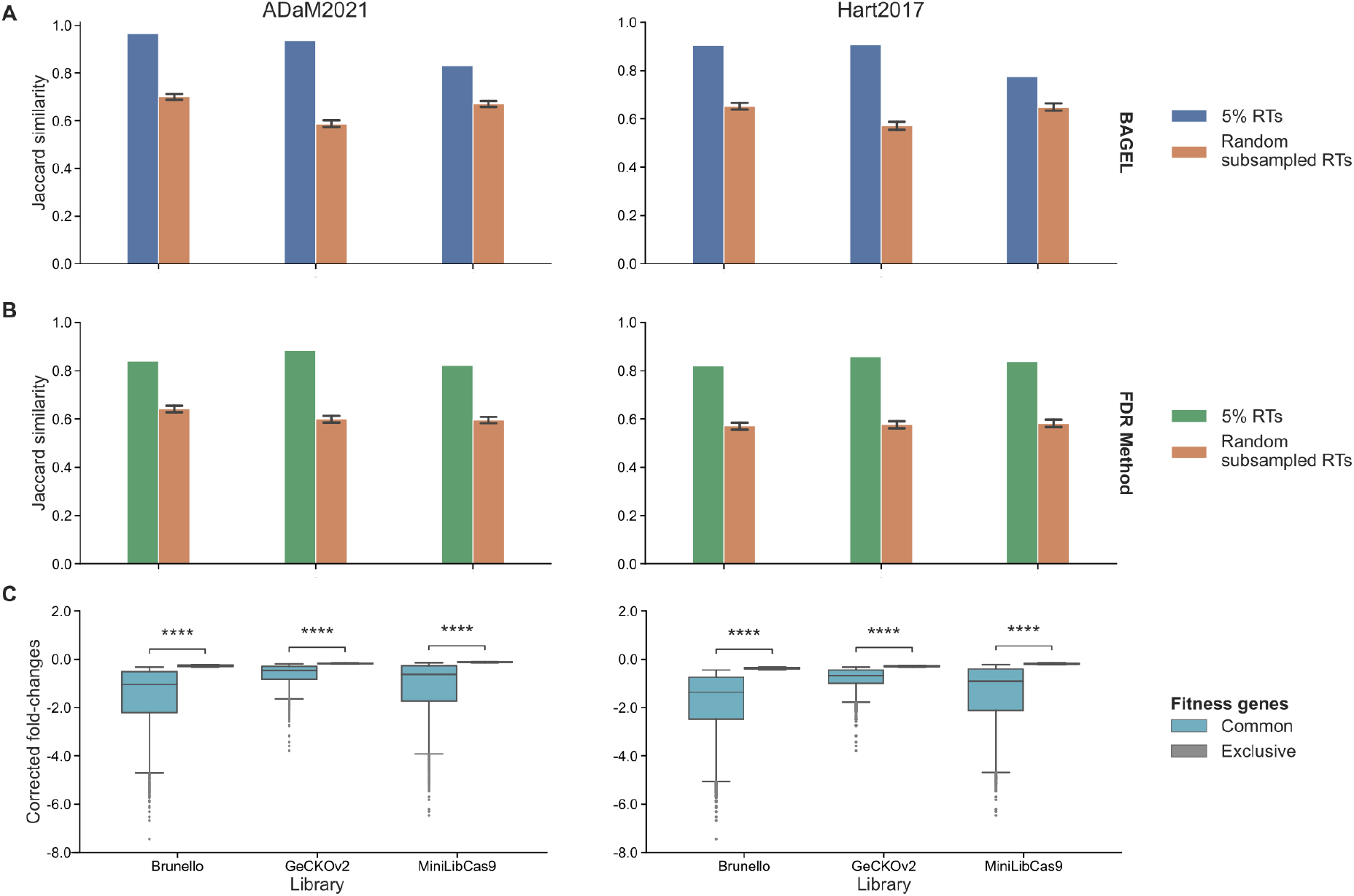
Validation of the MinTEs 5% library-independent reduced template (RT) on independent datasets. **AB**. Jaccard Similarity scores computed between fitness gene-sets predicted by BAGEL (**A**) or the FDR method (**B**), using as template classifier the MinTEs 5% library-independent RTs - derived from one of two state-of-the art reference gene-sets (different columns) - or 1,000 randomly subsampled RTs (as indicated by the different colours), and the original reference gene-set of derivation, respectively. BAGEL is applied to three independent screens of the HT-29 cell lines employing different libraries, as indicated by the different groups in each plot. **C**. Gene depletion fold-changes (post-correction for copy-number-bias) for genes that are commonly predicted as fitness by BAGEL when using as template classifier the MinTEs 5% library-independent RTs or the original reference gene-set of derivation, and for genes that are exclusive to one of the two predicted sets (as indicated by the different colours), **** indicates p-value < 1e-4.

#### Use case B: Estimation of gene essentiality using the FDR method

We sought to assess the performances of our MinTEs library-independent RTs when using them as a template classifier in the False Discovery Rate (FDR) method (**Fig. 1B**) by comparing, as for the previous use case, sets of genes predicted to be fitness at a 5% FDR when using the MinTEs RTs versus using whole reference gene-sets.

As for the previous case, overall tested MinTEs input parameters and percentages of subsampling, the MinTEs library-independent RTs outperformed random-subsampling as well as DepMap-driven RTs both on training and validation libraries (**Fig. S6**), for which we observed Jaccard similarity scores remarkably higher than 0.9 for the vast majority of tested subsampling percentages.

Zooming in on the MinTEs library-independent RTs of minimal size (5%) highlighted much higher Jaccard similarity scores with respect to randomly selected RTs (**Fig. 4B**). Finally, also in this case, the MinTEs library-independent RTs abundantly outperformed the DepMap-driven RTs (**Fig. S7**).

#### Top-essential genes are faithfully recapitulated by the MinTEs reduced templates

We compared fitness genes obtained by applying BAGEL to the validation libraries’ datasets using the original reference gene-sets versus the MinTEs library-independent RTs to evaluate the cell viability reduction effect elicited by CRISPR-Cas9 targeting genes that were consistently and inconsistently predicted, i.e. common fitness genes and exclusive fitness genes, respectively (**Fig. 4C**). For each independent CRISPR-Cas9 library we performed a Welch t-test between gene depletion FCs of common versus exclusive fitness genes. We observed that for the MinTEs RT of minimal size (5%) the common fitness gene FCs were systematically significantly lower (p < 1e-4), indicative of a stronger fitness reduction, than those of the exclusive genes. This implies (1) that the exclusive genes (missed by the MinTEs RTs) are more likely to be misclassified due to their borderline essentiality, and (2) that strong essential gene hits are always retained. Moreover, observing this for the RT of minimal size guarantees that this finding holds also for larger templates.

#### Use case C: Scaling gene depletion fold-changes

We evaluated the performances of the MinTEs RTs when used as a template to scale gene essentiality profiles derived from CRISPR-Cas9 screens. This is a common practice for inter-screen comparability and data interpretability (Aguirre et al. 2016; Munoz et al. 2016; Pacini et al. 2021; Dempster, Pacini, et al. 2019; Meyers et al. 2017; Tsherniak et al. 2017). This scaling employs FC distributions of reference EG and NEG to estimate the coefficients of a min-max normalisation following which the median depletion FCs of the EGs is equal to -1 and the median of the NEG is equal to 0 (**Fig. 1C**).

For this analysis, we considered the MinTEs training datasets (**Fig. S8**,**S9**), as well as an independently generated cancer dependency dataset derived from a compendium of three large-scale RNAi screenings covering 712 unique cancer cell lines, and processed with the DEMETER2 model (McFarland et al. 2018) (the DEMETER dataset, **Fig. 5A**). In addition, we focused on the assessment on the MinTEs library-independent RT of minimal size (5%), as larger percentages would perform at least as good (as shown in the previous sections (**Fig. 3**).

**Fig. 5.**
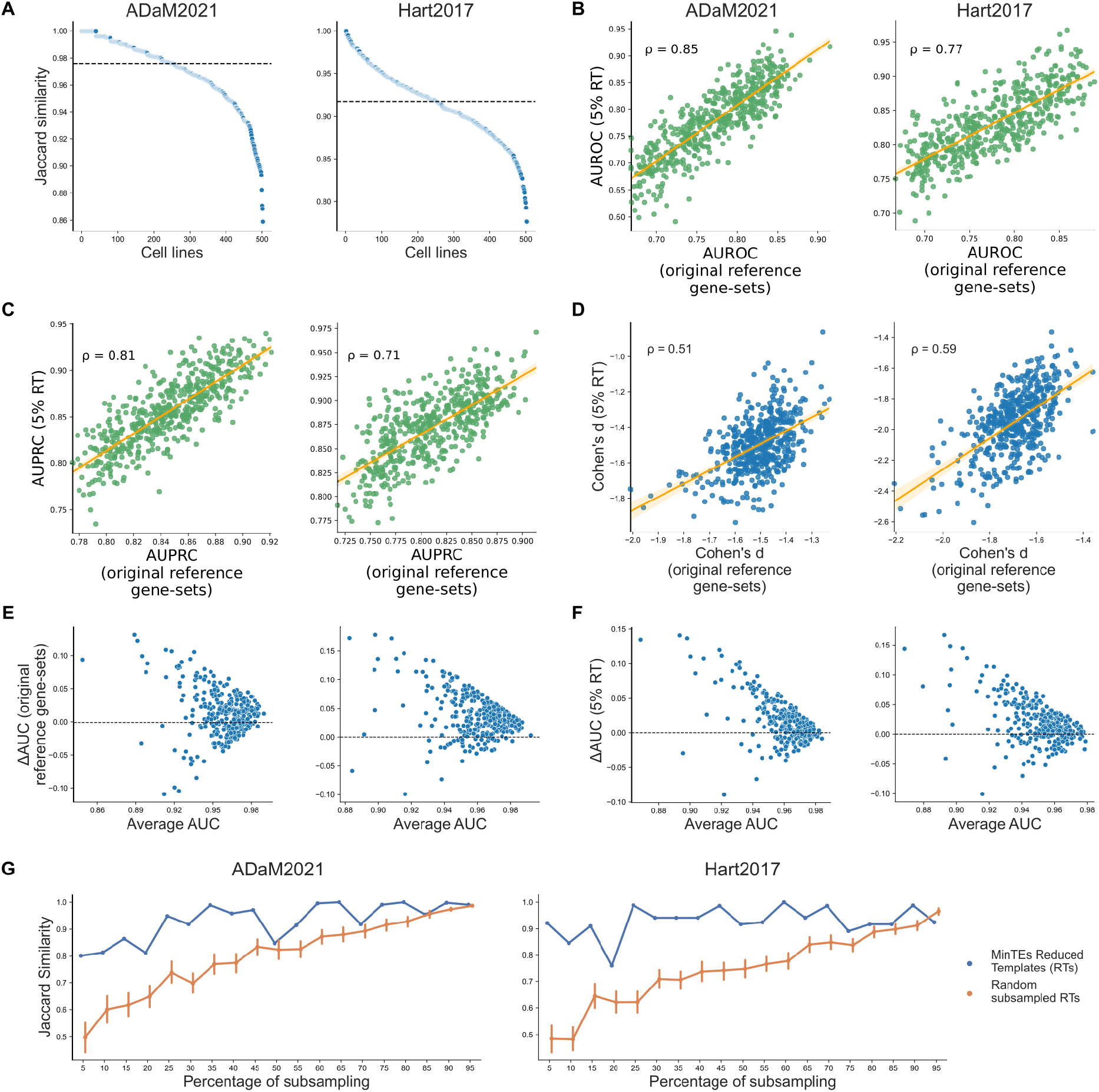
Depletion fold-change scaling and quality control assessment using MinTEs library-independent minimal templates. In all the plots, points indicate individual cell line screens, with different considered original reference essential gene-sets (ADaM2021 or Hart2017 gene-sets) indicated by different columns. **A**. Jaccard similarity scores between essential gene-sets called post scaling the DEMETER dataset with the whole reference gene-sets and the MinTEs 5% reduced templates (RTs), respectively. Dashed lines indicate median values. **BC**. Area Under Receiver Operating Characteristic (AUROC) **(B)** and Area Under Precision Recall Curve (AUPRC) **(C)** computed on the DEMETER dataset using the whole reference gene-sets (x-axis) or the MinTEs 5% RTs (y-axis), with Pearson’s correlation coefficients. **D**. Cohen’s d scores computed on the DEMETER dataset using the whole reference gene-sets (x-axis) or the MinTEs 5% RTs (y-axis), with Pearson’s correlation coefficients. **EF**. Bland-Altman plots comparing BAGEL and MAGeCK performances in terms of ability to correctly classify essential and nonessential genes. The two methods are applied to the project Score cancer dependency dataset. Values on the y-axis indicate the difference in performance between BAGEL and MAGeCK when applied to an individual cell line screen. Values on the x-axis indicate the average of the two AUC values, thus the overall screen quality. Plots in **(E)** are obtained using the original reference gene-sets as a template classifier for BAGEL, whereas plots in **(F)** are obtained using the MinTEs RTs of minimal size (5%). **G**. Average Jaccard Similarity scores, of fitness gene-sets obtained with BAGEL when using as classification templates the MinTEs derived RTs or randomly selected RTs (as indicated by the different colours) obtained from two state-of-the-art reference essential gene-sets (different columns) and the same reference set of non essential genes), on an organoid derived from colorectal patient sample, across percentage of subsampling. For random performances, the bars show 95% confidence interval computed across 1,000 train-test reshufflings of the input dataset for each combination of parameters (i.e. reference gene-set and percentage of subsampling).

Specifically, for each combination of dataset (KY/AVANA CRISPR-Cas9 or DEMETER) per initial reference gene-set (ADaM2021 or Hart2017), we scaled the dataset on a screened cell line basis using the whole reference gene-sets or the MinTEs 5% RT, then in each screen we call essential genes those with a scaled depletion FC lower than -0.5, i.e. equivalent to half the fitness effect of the EG used for the scaling (as done in other studies (Pacini et al. 2021; Dempster, Pacini, et al. 2019; Tsherniak et al. 2017)).

For every screened cell line, we focused on a set of prior known bona-fide essential genes independently assembled (from (Pacini et al. 2021; Subramanian et al. 2005; Iorio et al. 2018)), and determined which of these were called essential according to the scaling threshold, defining sets of recalled bona-fide essential (one for each cell line). Finally, we computed Jaccard scores between sets of recalled bona-fide essential obtained when scaling the dataset under consideration with the whole original reference gene-sets or the MinTEs 5% RTs, obtaining one score per screened cell line.

We observed very high Jaccard scores invariantly across datasets and starting reference gene-sets (median across cell lines > 0.9, and > 0.94 in 5 out of 6 cases) (**Fig. 5A**). Interestingly, the ADaM2021 derived MinTEs RTs outperformed the Hart2017 counterparts (minimum median Jaccard score > 0.97 against 0.90) across all instances (**Fig. 5A, S8**). In addition, we extended the analysis to all fitness genes with a scaled depletion FC lower than -0.5, and computed Jaccard similarity scores for each screened cell line as well. The ADaM2021 derived MinTEs RTs consistently showed higher median Jaccard score than the Hart2017 counterparts, with a larger gap when focusing on the DEMETER dataset (minimum median Jaccard score > 0.89 against 0.65) (**Fig. S9**), confirming a higher level of reliability for this collection of template EGs and NEGs (Vinceti et al. 2021).

#### Use case study D: Quality control assessment via ROC analysis

Common metrics used to assess the quality of CRISPR-Cas9 screens (Behan et al. 2019; Sanson et al. 2018; Dempster, Pacini, et al. 2019) are the areas under Receiver Operating Characteristics (AUROC) curve or the Area Under the Precision-Recall Curve (AUPRC) obtained when considering the gene FC profiles as rank based classifier of reference gene-sets of EGs and NEGs (**Fig 1D**). We computed AUROCs and AUPRCs for the gene FC profiles in the DEMETER dataset (as well as the MinTEs training dataset) using the reference gene-sets and the MinTEs RTs of minimal size (5%). These tested datasets contain only high-quality cell lines, extensively filtered following quality control (Behan et al. 2019; McFarland et al. 2018; Dempster, Rossen, et al. 2019). So to extend our benchmarking to a wider range of quality scores we pre-applied a noise injection strategy (STAR *Methods*). This noise-injection randomly moves the FC distributions of essential and nonessential original reference gene-sets distributions closer to each other in the individual cell line screens. This causes misclassification behaviours and allows to simulate low- and middle-quality screens.

We observed high correlations between AUROCs and AUPRCs computed when using MinTEs RTs versus whole reference sets across datasets and reference gene-sets of origin (R > 0.71, p-value below machine precision (< 10^−16^), **Fig. 5BC**) and (**Fig. S10**,**S11**). Particularly, we observed higher correlations on the training datasets (median R = 0.89 for the AUROC analysis, median R = 0.91 for the AUPRC analysis), with the ADaM2021 derived MinTEs RTs showing better performances than the Hart2017 counterparts.

#### Use case study E: Quality control assessment via Cohen’s d

Another common way to evaluate the quality of a CRISPR-Cas9 screen is by assessing the separation of the EG/NEG depletion fold-change distributions in terms of their mean values related to their pooled standard deviation, thus computing a screen overall effect size (**Fig 1E**). This can be quantified for example using the Cohen’s d coefficient (Cohen 2013; Behan et al. 2019). CRISPR-Cas9 screens with large negative scores having marked separation between the two reference sets’ FCs are of higher quality (STAR *Methods*).

We evaluated the performances of the MinTEs RTs when employed in this type of analysis. To this aim, we focused on the MinTEs training datasets (**Fig. S12**) as well as the DEMETER dataset (**Fig. 5D**), and considering again only the MinTEs RTs of minimal size (5%), as the worst case scenario.

We observed that Cohen’s d computed when using MinTEs RTs versus whole reference sets were significantly correlated invariantly across datasets and reference gene-sets of origin (R > 0.51, p-value below machine precision (< 10^−16^), **Fig. 5D**) and (**Fig. S12**). We observed higher correlations on the training datasets (median R = 0.89) with slightly variability towards high-quality screens.

#### MinTEs reduced templates preserve BAGEL performances

We performed an analysis to investigate to what extent the tendency of BAGEL to outperform MAGeCK (Li et al. 2014) on data from mid-to-low quality screens is preserved when using a MinTEs template of minimal size as template classifier instead of the whole reference gene-sets.

To this aim we processed the Project Score dataset (Dwane et al. 2021; Behan et al. 2019) with MAGeCK as well as with BAGEL using either the original reference gene-sets and the MinTEs 5% library-independent RT as template classifiers (**Fig. 5EF**, STAR *Methods*).

We then considered BAGEL gene bayesian-factors and MAGeCK gene depletion FDRs yielded by each individual cell line screen as a rank based classifier of the EGs and NEGs in the BAGEL reference gene template, and computed an Area under the ROC curve (AUC) to quantify corresponding classification performances.

We computed the average between BAGEL- and MAGeCK-AUC to estimate the overall quality of an individual screen and their difference as an indicator of differential performances between BAGEL and MAGeCK.

We observed that BAGEL performances were superior to those of MAGeCK in an inversely correlated fashion with respect to overall screen quality when using the original reference gene-sets. This tendency was preserved when using the MinTEs 5% RTs as classification template for BAGEL (**Fig. 5F**).

#### The library-independent reduced templates are not model specific

To assess the generality of our RTs and their suitability for the analysis of scale-limited screens performed on more complex models than cell lines, we assessed the performances of the MinTEs derived library-independent RTs against genome-wide CRISPR-Cas9 screens performed on three organoids (Gonçalves et al. 2021), one derived from colorectal carcinoma (**Fig. 5G**) and two derived from oesophageal cancer (**Fig. S13**).

First, we preprocessed the raw read counts available in the study to correct for copy-number bias and derived fold-changes (STAR *Methods*). Then we compared the Jaccard similarity scores as detailed in *Performance assessment of library-specific reduced templates*, i.e. quantifying the similarity between BAGEL outputted fitness gene-sets when used with the whole reference gene-sets versus library-independent RTs. We also tested randomly subsampled RTs, and performed 1,000 samplings per subsampling percentage.

The library-independent RTs consistently outperformed the random RTs across reference gene-sets, organoid sample and percentage of subsampling. These results are comparable, in terms of performances, to those obtained on the datasets generated from independent CRISPR libraries (**Fig. S4**) and demonstrate that even for screens performed on cancer organoids our RTs outperform random subsampling, thus are not specific to cancer cell lines.

## Discussion

The identification of fitness genes is a pivotal applications of CRISPR-Cas9 screens, allowing us to better elucidate the inner workings of human cells and to accelerate the identification of suitable oncotherapeutic targets. To facilitate different types of supervised computational analyses of scale-limited CRISPR-Cas9 screens (requiring large reference sets of control genes) we developed a computational framework that identifies optimised reduced templates (RTs) of reference essential and nonessential gene-sets.

Our strategy identifies RTs that outperform both a random selection (random RTs) and a data driven selection based on large cancer dependency map datasets (DepMap-driven RTs). The latter constitutes indeed an intuitive biologically-inspired approach to select RTs at different percentages of subsampling, by picking genes that are observed to be always/never-essential. Of note, DepMap-driven RTs utterly fail at recapitulating the fold-change distributions of the original reference gene-sets, with MinTEs RTs showing a substantial improvement over these sets, especially for small subsampling percentages.

Here we demonstrate that RTs can be reliably used across a range of tested analytical scenarios, reference gene-sets, CRISPR-Cas9 library and percentages of subsampling, as they provide results that are very similar to those obtained when using state-of-the-art reference sets of essential/non-essential genes (ADaM2021 and Hart2017), retaining the vast majority of top essential gene hits across multiple screens and libraries. Thus, these RTs allow a wide range of analyses of scale-limited CRISPR-Cas9 screens, while having a reduced impact on the size of the employed library.

Importantly, our analyses suggest that the ADaM2021 derived MinTEs RTs might be better suited than Hart2017 counterparts, as they generally offer higher and more stable performances, especially for the quality control and scaling of CRISPR-Cas9 screens.

Our RTs are particularly relevant given how the use of full-size reference gene-sets clashes with -- or sometimes renders entirely unfeasible -- the design of a scale-limited screen. Current CRISPR-Cas9 screens require a vast number of cells in order to avoid epistatic effects that might arise due to the entrance of multiple guides in the same cell (Olivieri and Durocher 2021). This hampers the application of supervised analyses (e.g. BAGEL) that leverage reference templates to identify essential genes in complex models, such as orgainoids, as well as *in vitro* screens focusing on a small set of genes, since they would require a high number of screened reference genes and a large volume of cells.

In addition, CRISPR libraries show discrepancies arising from different reagents’ performances (i.e. sgRNA on-target efficiency and off-target effects). For these reasons, library-specific RTs show better performances than library-independent RTs on the dataset they were generated from. Therefore, researchers planning to perform scale-limited screens using the KY/AVANA CRISPR libraries should opt for the library-specific templates.

Here, we present a reliable strategy to reduce reference gene-sets down to 5% of the state-of-the-art sets, while still preserving optimal performances. This strategy yielded sets of RTs that are also able to retain high performances on more complex models, such as organoids. Our methodology has the potential to reduce costs and save time for scale-limited screens and to simplify organoid and other complex screens, opening up opportunities for investigating gene essentiality in a variety of pathologically relevant contexts.

## Limitations of the study

While we have shown that the RTs are not model specific, validating them on organoids’ screens, we have not tested them on data from patient-derived xenograft or in vivo screens (due to lack of data). These constitute more complex models than organoids for which the performances of the library-independent RTs might be less optimal than those reported in this study. Another important factor to take into account is the possible disease specificity of our RTs. Indeed, we tested the RTs on a variety of publicly available datasets, but we focused our efforts only on cancer models. Therefore, the RTs might not be suited for investigating other contexts.

Finally, we developed a computational framework to assemble library-specific and independent RTs: better templates for other CRISPR libraries might be derived in the future when new data will be available. At the moment, we were able only to derive AVANA and KY-specific RTs due to the large number of *in vitro* models that were screened with these two libraries. Assessing the contribution of several CRISPR libraries might lead to the generation of more robust library-independent templates.

## STAR Methods

### Source of reference essential and nonessential gene-sets

We downloaded the Hart2017 reference gene-set of essential genes (EG) (Traver Hart et al. 2017) and Hart2014 reference gene-set of nonessential genes (NEG) (Traver Hart et al. 2014) from https://github.com/hart-lab/bagel. The second EG set (i.e. ADaM2021) is from (Vinceti et al. 2021), in which we derived it through the application of a semi-supervised algorithm, the Adaptive Daisy Model (ADaM), on the latest version of the integrated Sanger and Broad depletion fold-change (FC) matrix (Pacini et al. 2021), preprocessed with CERES (Meyers et al. 2017) and available from the DepMap portal (https://www.depmap.org/broad-sanger/integrated_Sanger_Broad_essentiality_matrices_20201201.zip).

### Genome-wide CRISPR-Cas9 datasets acquisition and pre-processing

MinTEs is trained on two genome-wide CRISPR-Cas9 screen datasets: the Project Score release 1, screened using the KY v1.1 library (Tzelepis et al. 2016; Behan et al. 2019), and the Achilles release 21Q2, screened using the AVANA library (Doench et al. 2016). For the Project Score dataset, the raw read count files were downloaded from the Project Score portal (https://score.depmap.sanger.ac.uk/downloads). As illustrated in (Behan et al. 2019), only cell line screens passing a low and high quality threshold (see Methods, https://www.nature.com/articles/s41586-019-1103-9) were considered for downstream analyses (325 cell lines).

The Achilles 21Q2 dataset was downloaded from the DepMap portal (https://depmap.org/portal/download/). In particular, for consistency in correcting sgRNAs counts with the same tool (CRISPRcleanR (Iorio et al. 2018)), we downloaded raw read counts that did not undergo CERES application (Meyers et al. 2017), together with guide file (Achilles_guide_map.csv, https://depmap.org/portal/download/), mapping genomic coordinates of sgRNAs to their target gene, and the replicate file (Achilles_replicate_map.csv, https://depmap.org/portal/download/), mapping replicate IDs to their cell name ID and including batch and quality control assessment as explained in (Dempster, Rossen, et al. 2019). We included in further analysis only replicates passing quality control.

Afterwards, we converted raw read counts to cell-wise sgRNA corrected FCs via CRISPRcleanR v2.2.1 (Iorio et al. 2018) coherently for both datasets in order to account for gene-independent cell responses. In this way, we obtained 325 and 859 cell lines, respectively, for Project Score and Achilles datasets respectively.

### Generation and prioritisation of library-specific minimal templates

MinTEs aims at finding the best reduce templates (RTs), for a fixed percentage of subsampling, reference EGs and chosen CRISPR-Cas9 library, that maximises the Jaccard similarity on the input dataset with respect to results that would be obtained with the original reference gene-sets. Particularly, MinTEs takes as input a CRISPR-Cas9 dataset of sgRNA-level depletion FCs and reference EG and NEG sets. The input dataset, containing CRISPRcleanR corrected sgRNA-level FCs, is first randomly split column-wise (i.e. cell-wise) into a training set and test set (typically following a 80/20 partition), then MinTEs initialises a random subset of EG_s_ and NEG_s_, whose size is defined by the percentage of subsampling *s*. BAGEL v115 (https://github.com/hart-lab/bagel) is then executed screen-wisely along the training set to compute, for each gene *g*, the Bayes Factor scores (BF_g_). For each cell line, we then ranked BF_g_ scores of the reference EG and NEG in increasing order. The ranked list is used to calculate positive predictive values (PPVs) as follows. For each rank position *i*, we calculated a set of predicted fitness genes (PFG) as:

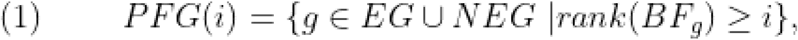

where *rank*(BF_g_) is the corresponding rank position of gene *g* in the reference gene set based on BF score. Thus, PPV in the *i*^*th*^ rank position is defined as:

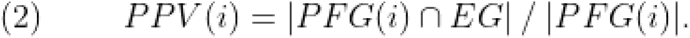

We determined the lowest threshold of BF score (BF*) in rank position *i\** such that PPV(*i*^*\**^) ≥ 0.95, which is equivalent to an FDR of 5%. The BF* threshold was calculated screen-wisely and then subtracted from the profile of BF_g_ scores. All genes having a BF_g_ score > 0 were classified as fitness genes, leading to a final set of rank independent predicted fitness genes (PFGs).

In each iteration of the optimisation process and for a CRISPR-Cas9 library *l* and a cell line *m*, the PFG sets derived from the reduced templates (RTs) 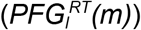 were then compared cell-wisely to those obtained by running BAGEL using the original reference gene-sets (*PFG*_*l*_*(m)*). The cost function to optimise was the Jaccard Similarity (JS), averaged across cells in the training set partition (*Tr*), defined as the size of the intersection divided by the size of the union of two sets:

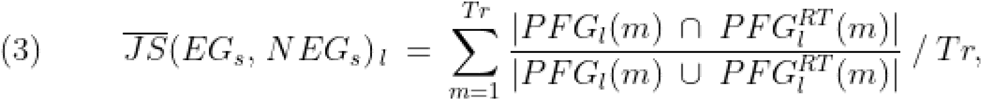

After the first iteration, the procedure was repeated by swapping genes in the EG_s_ and NEG_s_ sets with those in the hold-out sets, i.e. EG \ EG_s_ and NEG \ NEG_s_ respectively, according to the value of the substitution rate (*SR*) (see MinTEs hyperparameters optimisation for details). Jaccard similarity for the updated RT was then computed via (3) with MinTEs storing at each iteration the RT with the highest average JS. Defining with *C* hyperparameter the maximum number of iterations without improvements, the greedy search proceeds until the algorithm does not obtain any improvement (default is no improvement on the fourth decimal position) after C consecutive iterations or when SR leads to the base case (only one gene is swapped either in EG_s_ or NEG_s_ sets), meaning that is plateau is reached. The first scenario happens in the static instance of SR, the second one in the dynamic case (see MinTEs hyperparameters optimisation for details). Hence, MinTEs selects the best sets of EG_s_ and NEG_s_ that maximises the objective function in (4), namely

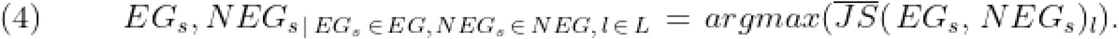

For each combination of parameters (i.e. CRISPR-Cas9 library *l*, EG set and percentage of subsampling), we carried out 1,000 train-test reshufflings. Once MinTEs reaches termination on the training set, it computes the average JS using equation (3) but for the cell left in the testing set and having EG_s_ and NEG_s_ as obtained from (4). As a baseline, we run MinTEs with a static SR equal to 1 and C set to 0, for each combination of parameters. This is the same as performing a one-off random subsampling of EG_s_ and NEG_s_. In the case of random subsampling, the average JS was computed on the total number of cell lines without splitting in training and test sets.

Hence, via the reshuffling procedure, MinTEs delivered 1,000 optimised RTs and 1,000 random RTs for each combination of parameters. Finally, we devised a prioritisation strategy to select the best library-specific RTs for each combination of parameters by applying the following formula:

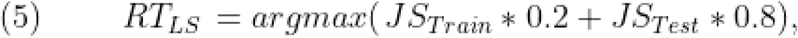

where JS_Train_ is the average score obtained from the library-specific RT set on the training set, and JS_Test_ is the average score on the test set. We mainly rewarded the performances obtained on the test set in order to favour RT sets with higher performances on never-before-seen data, and picked the RT with the highest library-specific score.

### Prioritisation of library-independent minimal templates

We implemented an additional prioritisation strategy to select the top-performing RTs in a library-independent manner. First, we cross-tested each pair of RT sets outputted by MinTEs on their respective never-before-seen CRISPR-Cas9 dataset (i.e. KY-specific RT sets were tested on the Achilles dataset, obtained with the AVANA library, and vice versa), according to the EG set and percentage of subsampling. Then, we implemented a similar prioritisation strategy as for selection of the library-independent RTs, following the formula:

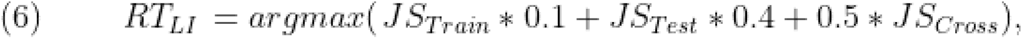

where JS_Train_ is the average score obtained from the gene subsets on the training set, JS_Test_ is the average score on the test set, and JS_Cross_ is the average score on the other CRISPR-Cas9 dataset. Here, we mainly rewarded the performances on the never-before-seen CRISPR-Cas9 datasets and the test set in the dataset used during the training, and selected the RT with the highest library-independent score.

### MinTEs hyperparameters optimisation

MinTEs has two hyperparameters: the substitution rate (*SR*), defined as the fraction of genes swapped at each iteration between the RT and the hold-out set, and *C*, defined as the maximum number of iterations without improvements in terms of Jaccard Similarity. For tuning the hyperparameters, we tested MinTEs on the Project Score release 1 dataset (https://score.depmap.sanger.ac.uk/downloads) at 5% subsampling using 80% of the cell lines as training and the remainder as test set, and performing 1,000 train-test reshufflings. Here, we selected the smallest percentage of subsampling for the hyperparameters fine-tuning as changes in this setting lead to larger differences in performances.

First, we tested “static” and “dynamic” instances of SR. In the static setting, a fixed number of genes is swapped at each optimisation step and MinTEs stops after reaching C steps with no improvement in performance (default is no improvement on the fourth decimal position). We tested the following values of static *SR* settings: 0.05, 0.15, 0.25, 0.5, 0.75 and 1. For instance, let’s consider 5% as a subsampling percentage, a EG set of 684 genes, a NEG set of 927 genes and a static SR of 0.25:

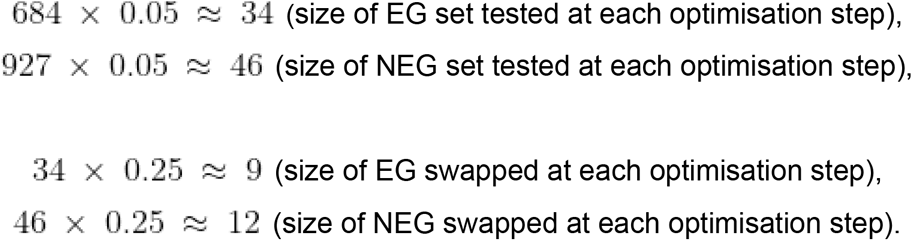

According to the example, MinTEs randomly swaps 9 EG and 12 NEG in the RT derived from the previous iteration with the same number of genes sampled from the hold-out set (in the first step the subsets are randomly initialised). Then, the algorithm compares the original and the newly-sampled RT, and selects that having the higher average Jaccard score.

On the other hand, in the dynamic setting for a given percentage of subsampling *s*, SR follows an exponential decay function as shown in formula (7) below:

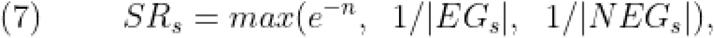

where *e* is the Nepero constant, |*EG*_*s*_| and |*NEG*_*s*_| are the sizes of the EG and NEG sets at percentage of subsampling *s*. The exponent *n* is initialised at 0 leading to a full random replacement at the beginning, it is then constantly increased by one after C consecutive iterations without improvement (default is no improvement on the fourth decimal position). A decrease in SR implies that a lower number of genes are swapped by MinTEs, until it reaches the base case (i.e. only one gene either in the EG_s_ or the NEG_s_ sets are swapped). For testing, C was set to 10 for all instances of SR (both static and dynamic) in order to avoid overfitting. For static SR, MinTEs stopped after 10 consecutive iterations without improvement, whereas for the dynamic setting, it continued until reaching the base case. The dynamic SR showed better performances (**Fig. 2B**), allowing us to explore combinations of subsets that otherwise would not have been accessible using a static SR. Thus, it was used as the default setting for the algorithm.

In order to find the optimal value for C, we tested all values 5 ≤ x ≤ 200, with a constant increment of 5. Again, MinTEs was run on the Project Score dataset at 5% subsampling and using the dynamic SR. We concluded 20 to be the optimal value for C (**Fig. 2C**), constituting the best trade-off between performance on the test sets and computational cost of the optimisation procedure.

### Performance assessment of library-specific reduced templates

We tested the library-specific RTs on the training datasets (Project Score release 1 with KY library, and Achilles 21Q2 with AVANA library) using BAGEL for the estimation of gene essentiality. We computed the average Jaccard score by comparing the predicted fitness genes from the whole original reference gene-sets and library-specific RTs for every combination of parameters. We compared the performances of the library-specific RT against random as well as DepMap-driven RTs. Random RTs were randomly drawn, whereas the DepMap-driven RTs were obtained by selecting EGs and NEGs with lowest and highest average FCs respectively, and in the same number of library-specific RTs.

### Validation of library-independent reduced templates

#### Estimation of gene essentiality using BAGEL or the FDR method

We validated the library-independent RTs on the training datasets (i.e. Project Score release 1 and Achilles 21Q2), as well as three independent datasets obtained with orthogonal CRISPR-Cas9 libraries derived from the following genome-wide CRISPR-Cas9 libraries: Brunello (Sanson et al. 2018), GeCKOv2 (Sanson et al. 2018) and MiniLibCas9 (Gonçalves et al. 2021). All three screens were performed on the HT-29 cell line. We preprocessed the raw count files following the same pipeline used as for the training datasets, generating corrected sgRNA-level FCs. We then applied BAGEL using the whole reference gene-sets or library-independent RTs as template classifiers, computed fitness genes at 5% FDR, and compared the performances in terms of Jaccard scores, for every combination of parameters (use case study A).

We carried out the same procedure for the False Discovery Rate (FDR) method, but predicted fitness genes were inferred directly from gene-level FC datasets as opposed to Bayes factors (use case study B). For the training datasets, we compared the performances of the library-independent RTs against random and DepMap-driven RTs for both gene essentiality estimation methods. Instead, for the validation datasets, we compared the performances of library-independent RTs only against those of the random RTs.

#### Quality control assessment and data scaling of CRISPR-Cas9 screens

We also validated the library-independent RTs based on widely adopted strategies to perform data normalisation and assess CRISPR-Cas9 screen quality control. To this aim, we considered the training datasets (Achilles 21Q2 and Project Score release 1) as well as an integrated dependency dataset derived from the aggregation of three large-scale RNAi screening datasets after applying the DEMETER2 model (https://github.com/cancerdatasci/demeter2). DEMETER dataset covers a total of 712 unique cancer cell lines (available at https://ndownloader.figshare.com/files/11489669).

For the use case study C, we performed a scaling of the MinTEs training datasets and the DEMETER dataset, using the original reference gene-sets and the 5% library-independent RTs. The scaling was performed cell-wise using the function CoRe.scale_to_essentials from the CoRe R package v1.0.2 (https://github.com/DepMap-Analytics/CoRe). Then, we assembled a set of bona-fide reference essential genes independently derived from (Pacini et al. 2021; Subramanian et al. 2005; Iorio et al. 2018), defined as essential those with a FC < -0.5, and computed the Jaccard score between the original reference gene-sets and 5% library-independent RTs. We carried out the same analysis considering all the fitness genes with FC < -0.5 as well, and computed cell-wise Jaccard similarity scores.

For the use case study D, we assessed screen quality using two common metrics, namely the Area Under Receiver Operating Characteristic (AUROC) and the Area Under Precision Recall Curve (AUPRC). We applied a noise injection strategy to move closer the FC distributions of the original reference gene-sets, in order to simulate low-to middle-quality screens. For each cell line, we first computed the distance *d* between the original EG and NEG’ medians, then we randomly added to EG’ FCs and subtracted to NEG’ FCs a corrective factor in the range of [0 ·d, 0.25 · d]. Afterwards, we computed cell-wise AUROCs and AUPRCs based on the processed EG and NEG’ FC distributions as true positive and negative controls, respectively. Finally, we compared the original reference gene-sets and 5% library-independent RT results in terms of Pearson correlation.

For the use case study E, we computed cell-wise Cohen’s d coefficients to measure the magnitude of separation between EG and NEG gene-sets’ FC distributions, using the original reference gene-sets and 5% library-independent RTs, and we compared the performances via Pearson correlation.

#### MinTEs reduced templates preserve BAGEL performances

Project Score release 1 dataset was used to compare BAGEL and MAGeCK (v 0.5.3, https://sourceforge.net/projects/mageck/) methods for the estimation of gene essentiality. For BAGEL, we used the original reference gene-sets and 5% library-independent RTs, with respect to the reference EG set, as template classifiers leading to four different datasets of Bayes Factor, and scaled them cell-wise at 5% FDR. For MAGeCK instead, we downloaded the dataset already processed with the method from the Project Score portal (https://score.depmap.sanger.ac.uk/downloads).

Then, we assessed the performances in terms of AUROC for every cell line of the datasets, processed either with BAGEL or MAGeCK. Finally, we computed the difference and average AUROC scores for each cell line between BAGEL- and MAGeCK-derived datasets.

#### The library-independent reduced templates are not model specific

The raw read counts of the organoids were preprocessed using CRISPRcleanR to correct for copy-number bias and generate corrected sgRNA-level FCs. We then reiterated the same procedure illustrated in the use case study A, and compared the Jaccard similarity scores, in terms of fitness genes estimated using BAGEL, between library-independent and random RTs compared to the whole reference templates.

## Supporting information

Revised Supplementary material

## Author contributions

AV conceived the study, designed, and performed the benchmark analyses, assembled the jupyter notebook, wrote and revised the manuscript. UP contributed to the design of the benchmark analyses and revised the manuscript. LT contributed to results’ interpretation, manuscript writing and revision. FI conceived the study, contributed to the design of the benchmark analyses, wrote, and revised the manuscript, supervised the study.

## Code availability

The code illustrating MinTEs computational framework is publicly available at https://github.com/AleVin1995/MinTEs. All results and figures can be reproduced on the fly at https://colab.research.google.com/github/AleVin1995/MinTEs/blob/master/notebooks/Summary_Colab.ipynb.

## References

Aguirre, Andrew J., Robin M. Meyers, Barbara A. Weir, Francisca Vazquez, Cheng-Zhong Zhang, Uri Ben-David, April Cook, et al. 2016. “Genomic Copy Number Dictates a Gene-Independent Cell Response to CRISPR/Cas9 Targeting.” Cancer Discovery 6 (8): 914–29.

Barrangou, Rodolphe. 2020. “Ushering in the Next CRISPR Decade.” The CRISPR Journal 3 (1): 2.

Behan, Fiona M., Francesco Iorio, Gabriele Picco, Emanuel Gonçalves, Charlotte M. Beaver, Giorgia Migliardi, Rita Santos, et al. 2019. “Prioritization of Cancer Therapeutic Targets Using CRISPR–Cas9 Screens.” Nature 568 (7753): 511–16.

Birsoy, Kivanç, Tim Wang, Walter W. Chen, Elizaveta Freinkman, Monther Abu-Remaileh, and David M. Sabatini. 2015. “An Essential Role of the Mitochondrial Electron Transport Chain in Cell Proliferation Is to Enable Aspartate Synthesis.” Cell 162 (3): 540–51.

Cohen, Jacob. 2013. Statistical Power Analysis for the Behavioral Sciences. 2nd ed. London, England: Routledge. https://doi.org/10.4324/9780203771587.

Condon, Kendall J., Jose M. Orozco, Charles H. Adelmann, Jessica B. Spinelli, Pim W. van der Helm, Justin M. Roberts, Tenzin Kunchok, and David M. Sabatini. 2021. “Genome-Wide CRISPR Screens Reveal Multitiered Mechanisms through Which mTORC1 Senses Mitochondrial Dysfunction.” Proceedings of the National Academy of Sciences of the United States of America 118 (4). https://doi.org/10.1073/pnas.2022120118.

Dempster, Joshua M., Clare Pacini, Sasha Pantel, Fiona M. Behan, Thomas Green, John Krill-Burger, Charlotte M. Beaver, et al. 2019. “Agreement between Two Large Pan-Cancer CRISPR-Cas9 Gene Dependency Data Sets.” Nature Communications 10 (1): 5817.

Dempster, Joshua M., Jordan Rossen, Mariya Kazachkova, Joshua Pan, Guillaume Kugener, David E. Root, and Aviad Tsherniak. 2019. “Extracting Biological Insights from the Project Achilles Genome-Scale CRISPR Screens in Cancer Cell Lines.” bioRxiv. https://doi.org/10.1101/720243.

DeWeirdt, Peter C., Annabel K. Sangree, Ruth E. Hanna, Kendall R. Sanson, Mudra Hegde, Christine Strand, Nicole S. Persky, and John G. Doench. 2020. “Genetic Screens in Isogenic Mammalian Cell Lines without Single Cell Cloning.” Nature Communications 11 (1): 752.

Doench, John G. 2018. “Am I Ready for CRISPR? A User’s Guide to Genetic Screens.” Nature Reviews. Genetics 19 (2): 67–80.

Doench, John G., Nicolo Fusi, Meagan Sullender, Mudra Hegde, Emma W. Vaimberg, Katherine F. Donovan, Ian Smith, et al. 2016. “Optimized sgRNA Design to Maximize Activity and Minimize off-Target Effects of CRISPR-Cas9.” Nature Biotechnology 34 (2): 184–91.

Dwane, Lisa, Fiona M. Behan, Emanuel Gonçalves, Howard Lightfoot, Wanjuan Yang, Dieudonne van der Meer, Rebecca Shepherd, Miguel Pignatelli, Francesco Iorio, and Mathew J. Garnett. 2021. “Project Score Database: A Resource for Investigating Cancer Cell Dependencies and Prioritizing Therapeutic Targets.” Nucleic Acids Research 49 (D1): D1365–72.

Girardi, Enrico, Adrián César-Razquin, Sabrina Lindinger, Konstantinos Papakostas, Justyna Konecka, Jennifer Hemmerich, Stefanie Kickinger, et al. 2020. “A Widespread Role for SLC Transmembrane Transporters in Resistance to Cytotoxic Drugs.” Nature Chemical Biology 16 (4): 469–78.

Gonçalves, Emanuel, Mark Thomas, Fiona M. Behan, Gabriele Picco, Clare Pacini, Felicity Allen, Alessandro Vinceti, et al. 2021. “Minimal Genome-Wide Human CRISPR-Cas9 Library.” Genome Biology 22 (1): 40.

Hartenian, Ella, and John G. Doench. 2015. “Genetic Screens and Functional Genomics Using CRISPR/Cas9 Technology.” The FEBS Journal 282 (8): 1383–93.

Hart, T., M. Chandrashekhar, M. Aregger, Z. Steinhart, K. R. Brown, and G. MacLeod. 2015. “High-Resolution CRISPR Screens Reveal Fitness Genes and Genotype-Specific Cancer Liabilities.” Cell 163. https://doi.org/10.1016/j.cell.2015.11.015.

Hart, Traver, Kevin R. Brown, Fabrice Sircoulomb, Robert Rottapel, and Jason Moffat. 2014. “Measuring Error Rates in Genomic Perturbation Screens: Gold Standards for Human Functional Genomics.” Molecular Systems Biology 10 (July): 733.

Hart, Traver, and Jason Moffat. 2016. “BAGEL: A Computational Framework for Identifying Essential Genes from Pooled Library Screens.” BMC Bioinformatics 17 (April): 164.

Hart, Traver, Amy Hin Yan Tong, Katie Chan, Jolanda Van Leeuwen, Ashwin Seetharaman, Michael Aregger, Megha Chandrashekhar, et al. 2017. “Evaluation and Design of Genome-Wide CRISPR/SpCas9 Knockout Screens.” G3 7 (8): 2719–27.

Iorio, Francesco, Fiona M. Behan, Emanuel Gonçalves, Shriram G. Bhosle, Elisabeth Chen, Rebecca Shepherd, Charlotte Beaver, et al. 2018. “Unsupervised Correction of Gene-Independent Cell Responses to CRISPR-Cas9 Targeting.” BMC Genomics 19 (1): 604.

Kim, Eiru, and Traver Hart. 2021. “Improved Analysis of CRISPR Fitness Screens and Reduced off-Target Effects with the BAGEL2 Gene Essentiality Classifier.” Genome Medicine 13 (1): 2.

Koike-Yusa, H., Y. Li, E. -. P. Tan, M. D. C. Velasco-Herrera, and K. Yusa. 2014. “Genome-Wide Recessive Genetic Screening in Mammalian Cells with a Lentiviral CRISPR-Guide RNA Library.” Nature Biotechnology 32. https://doi.org/10.1038/nbt.2800.

Li, Wei, Johannes Köster, Han Xu, Chen-Hao Chen, Tengfei Xiao, Jun S. Liu, Myles Brown, and X. Shirley Liu. 2015. “Quality Control, Modeling, and Visualization of CRISPR Screens with MAGeCK-VISPR.” Genome Biology 16 (December): 281.

Li, Wei, Han Xu, Tengfei Xiao, L. Cong Michael I. Love, Feng Zhang, Rafael A. Irizarry, Jun S. Liu, Myles Brown, and X. Shirley Liu. 2014. “MAGeCK Enables Robust Identification of Essential Genes from Genome-Scale CRISPR/Cas9 Knockout Screens.” Genome Biology 15 (12): p554.

McFarland, James M., Zandra V. Ho, Guillaume Kugener, Joshua M. Dempster, Phillip G. Montgomery, Jordan G. Bryan, John M. Krill-Burger, et al. 2018. “Improved Estimation of Cancer Dependencies from Large-Scale RNAi Screens Using Model-Based Normalization and Data Integration.” Nature Communications 9 (1): 4610.

Meyers, Robin M., Jordan G. Bryan, James M. McFarland, Barbara A. Weir, Ann E. Sizemore, Han Xu, Neekesh V. Dharia, et al. 2017. “Computational Correction of Copy Number Effect Improves Specificity of CRISPR-Cas9 Essentiality Screens in Cancer Cells.” Nature Genetics 49 (12): 1779–84.

Miles, Linde A., Ralph J. Garippa, and John T. Poirier. 2016. “Design, Execution, and Analysis of Pooled in Vitro CRISPR/Cas9 Screens.” The FEBS Journal 283 (17): 3170–80.

Munoz, Diana M., Pamela J. Cassiani, Li Li, Eric Billy, Joshua M. Korn, Michael D. Jones, Javad Golji, et al. 2016. “CRISPR Screens Provide a Comprehensive Assessment of Cancer Vulnerabilities but Generate False-Positive Hits for Highly Amplified Genomic Regions.” Cancer Discovery 6 (8): 900–913.

Olivieri, Michele, and Daniel Durocher. 2021. “Genome-Scale Chemogenomic CRISPR Screens in Human Cells Using the TKOv3 Library.” STAR Protocols 2 (1): 100321.

Pacini, Clare, Joshua M. Dempster, Isabella Boyle, Emanuel Gonçalves, Hanna Najgebauer, Emre Karakoc, Dieudonne van der Meer, et al. 2021. “Integrated Cross-Study Datasets of Genetic Dependencies in Cancer.” Nature Communications 12 (1): 1661.

Parrish, Phoebe C. R., James D. Thomas, Austin M. Gabel, Shriya Kamlapurkar, Robert K. Bradley, and Alice H. Berger. 2021. “Discovery of Synthetic Lethal and Tumor Suppressor Paralog Pairs in the Human Genome.” Cell Reports 36 (9): 109597.

Peets, Elin Madli, Luca Crepaldi, Yan Zhou, Felicity Allen, Rasa Elmentaite, Guillaume Noell, Gemma Turner, Vivek Iyer, and Leopold Parts. 2019. “Minimized Double Guide RNA Libraries Enable Scale-Limited CRISPR/Cas9 Screens.” bioRxiv. https://doi.org/10.1101/859652.

Roesch, Ferdinand, Molly OhAinle, and Michael Emerman. 2018. “A CRISPR Screen for Factors Regulating SAMHD1 Degradation Identifies IFITMs as Potent Inhibitors of Lentiviral Particle Delivery.” Retrovirology 15 (1): 26.

Sanson, Kendall R., Ruth E. Hanna, Mudra Hegde, Katherine F. Donovan, Christine Strand, Meagan E. Sullender, Emma W. Vaimberg, et al. 2018. “Optimized Libraries for CRISPR-Cas9 Genetic Screens with Multiple Modalities.” Nature Communications 9 (1): 5416.

Shalem, Ophir, Neville E. Sanjana, Ella Hartenian, Xi Shi, David A. Scott, Tarjei Mikkelson, Dirk Heckl, et al. 2014. “Genome-Scale CRISPR-Cas9 Knockout Screening in Human Cells.” Science 343 (6166): 84–87.

Shalem, Ophir, Neville E. Sanjana, and Feng Zhang. 2015. “High-Throughput Functional Genomics Using CRISPR-Cas9.” Nature Reviews. Genetics 16 (5): 299–311.

Slabicki, Mikolaj, Zuzanna Kozicka, Georg Petzold, Yen-Der Li, Manisha Manojkumar, Richard D. Bunker, Katherine A. Donovan, et al. 2020. “The CDK Inhibitor CR8 Acts as a Molecular Glue Degrader That Depletes Cyclin K.” Nature 585 (7824): 293–97.

Steinhart, Zachary, Zvezdan Pavlovic, Megha Chandrashekhar, Traver Hart, Xiaowei Wang, Xiaoyu Zhang, Mélanie Robitaille, et al. 2017. “Genome-Wide CRISPR Screens Reveal a Wnt-FZD5 Signaling Circuit as a Druggable Vulnerability of RNF43-Mutant Pancreatic Tumors.” Nature Medicine 23 (1): 60–68.

Subramanian, Aravind, Pablo Tamayo, Vamsi K. Mootha, Sayan Mukherjee, Benjamin L. Ebert, Michael A. Gillette, Amanda Paulovich, et al. 2005. “Gene Set Enrichment Analysis: A Knowledge-Based Approach for Interpreting Genome-Wide Expression Profiles.” Proceedings of the National Academy of Sciences of the United States of America 102 (43): 15545–50.

Su, Dan, Xu Feng, Medina Colic, Yunfei Wang, Chunchao Zhang, Chao Wang, Mengfan Tang, Traver Hart, and Junjie Chen. 2020. “CRISPR/CAS9-Based DNA Damage Response Screens Reveal Gene-Drug Interactions.” DNA Repair 87 (March): 102803.

Tarumoto, Yusuke, Bin Lu, Tim D. D. Somerville, Yu-Han Huang, Joseph P. Milazzo, Xiaoli S. Wu, Olaf Klingbeil, Osama El Demerdash, Junwei Shi, and Christopher R. Vakoc. 2018. “LKB1, Salt-Inducible Kinases, and MEF2C Are Linked Dependencies in Acute Myeloid Leukemia.” Molecular Cell 69 (6): 1017–27.e6.

Tsherniak, Aviad, Francisca Vazquez, Phil G. Montgomery, Barbara A. Weir, Gregory Kryukov, Glenn S. Cowley, Stanley Gill, et al. 2017. “Defining a Cancer Dependency Map.” Cell 170 (3): 564–76.e16.

Turner, David J., and Martin Turner. 2021. “RNA Binding Proteins As Regulators of Oxidative Stress Identified by a Targeted CRISPR-Cas9 Single Guide RNA Library.” The CRISPR Journal 4 (3): 427–37.

Tzelepis, Konstantinos, Hiroko Koike-Yusa, Etienne De Braekeleer, Yilong Li, Emmanouil Metzakopian, Oliver M. Dovey, Annalisa Mupo, et al. 2016. “A CRISPR Dropout Screen Identifies Genetic Vulnerabilities and Therapeutic Targets in Acute Myeloid Leukemia.” Cell Reports 17 (4): 1193–1205.

Vinceti, Alessandro, Emre Karakoc, Clare Pacini, Umberto Perron, Riccardo Roberto De Lucia, Mathew J. Garnett, and Francesco Iorio. 2021. “CoRe: A Robustly Benchmarked R Package for Identifying Core-Fitness Genes in Genome-Wide Pooled CRISPR-Cas9 Screens.” bioRxiv. https://doi.org/10.1101/2021.05.25.445610.

Wheeler, Emily C., Anthony Q. Vu, Jaclyn M. Einstein, Matthew DiSalvo, Noorsher Ahmed, Eric L. Van Nostrand, Alexander A. Shishkin, Wenhao Jin, Nancy L. Allbritton, and Gene W. Yeo. 2020. “Pooled CRISPR Screens with Imaging on Microraft Arrays Reveals Stress Granule-Regulatory Factors.” Nature Methods 17 (6): 636–42.

Williams, Robert T., Rohiverth Guarecuco, Leah A. Gates, Douglas Barrows, Maria C. Passarelli, Bryce Carey, Lou Baudrier, et al. 2020. “ZBTB1 Regulates Asparagine Synthesis and Leukemia Cell Response to L-Asparaginase.” Cell Metabolism 31 (4): 852–61.e6.

Zhang, Hao, Yang Zhang, Xinyue Zhou, Shaela Wright, Judith Hyle, Lianzhong Zhao, Jie An, et al. 2020. “Functional Interrogation of HOXA9 Regulome in MLLr Leukemia via Reporter-Based CRISPR/Cas9 Screen.” eLife 9 (October). https://doi.org/10.7554/eLife.57858.

Zhou, Yuexin, Shiyou Zhu, Changzu Cai, Pengfei Yuan, Chunmei Li, Yanyi Huang, and Wensheng Wei. 2014. “High-Throughput Screening of a CRISPR/Cas9 Library for Functional Genomics in Human Cells.” Nature 509 (7501): 487–91.

Zhu, Yuqi, Theodore Groth, Anju Kelkar, Yusen Zhou, and Sriram Neelamegham. 2021. “A GlycoGene CRISPR-Cas9 Lentiviral Library to Study Lectin Binding and Human Glycan Biosynthesis Pathways.” Glycobiology 31 (3): 173–80.

