## Supplementary material for "Reduced gene templates for supervised analysis of scale-limited CRISPR-Cas9 fitness screens": Revised Supplementary material

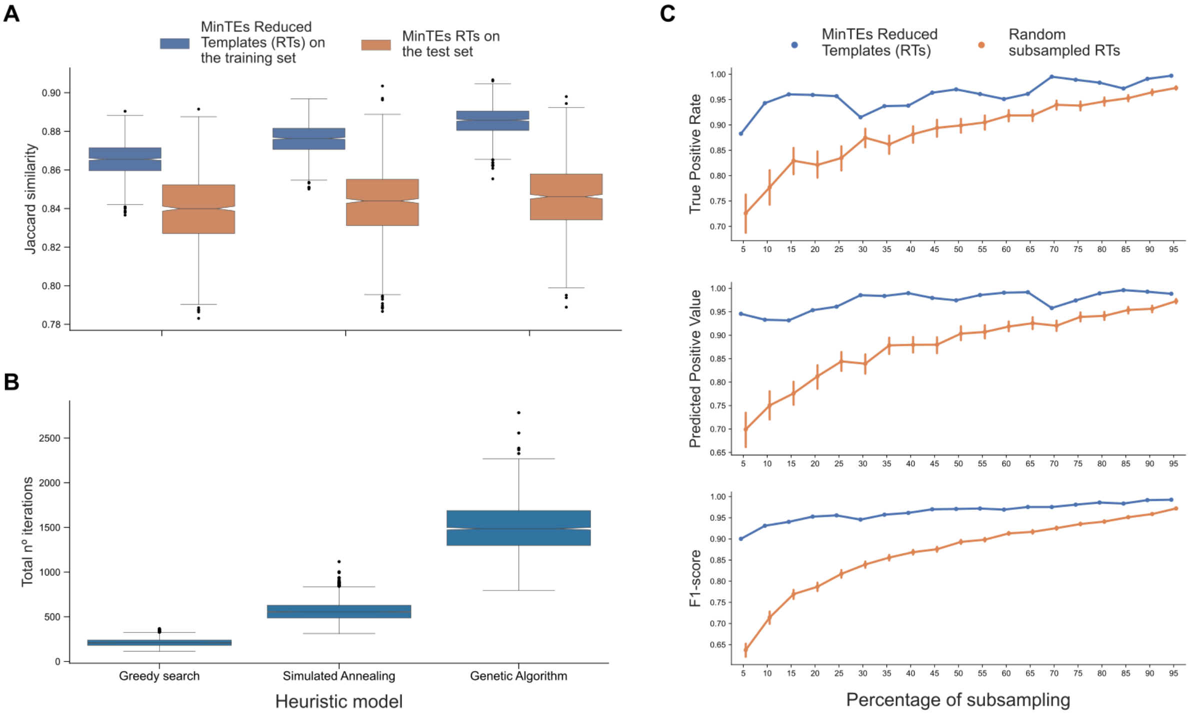


**Fig. S1 - Testing of different setups for MinTEs computational framework. A.** Average Jaccard Similarity scores of fitness gene-sets obtained with BAGEL when using as classification templates the MinTEs derived RTs, obtained using the Hart2017 essential and Hart2014 nonessential gene-sets on the Project Score dataset (obtained with the KY library) at 5% percentage of subsampling via different methods, i.e. greedy search, simulated annealing and genetic algorithm (x-axis). RTs' performances are split between training and test sets (different colours) and we performed 1000 train-test reshufflings. **B.** Total number of iterations needed by each method (x-axis) to achieve convergence. The models were tested using the Hart2017 essential and Hart2014 nonessential gene-sets on the Project Score dataset (obtained with the KY library) to identify RTs at 5% percentage of subsampling. **C.** Average True Positive Rate, Predicted Positive Value, and F1-score between fitness gene-sets predicted by BAGEL when using as classification templates the MinTEs derived library-specific RTs or randomly selected RTs (as indicated by the different colours) obtained from the Hart2017 reference essential gene-set and the Hart2014 reference set of nonessential genes, for the KY single guide RNA library, across percentage of subsampling. For random performances, the bars show 95% confidence interval computed across 1,000 train-test reshufflings of the input dataset for each combination of parameters (i.e. percentage of subsampling).

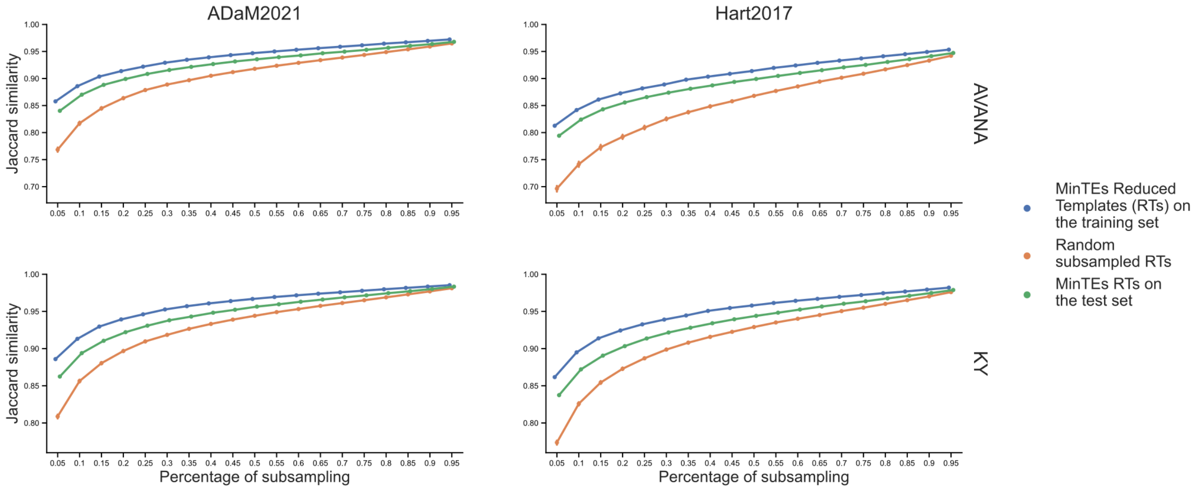


**Fig. S2 - Performances of MinTEs reduced templates (RTs).** Average Jaccard Similarity scores, of fitness gene-sets obtained with BAGEL when using as classification templates the MinTEs derived RTs or randomly selected RTs (as indicated by the different colours) obtained from two state-of-the-art reference essential gene-sets (different columns) and the same reference set of non essential genes), for two different single guide RNA libraries (different rows), across percentage of subsampling. For random and MinTEs performances, the bars show 95% confidence interval computed across 1,000 train-test reshufflings of the input dataset for each combination of parameters (i.e. reference gene-set, CRISPR-Cas9 library and percentage of subsampling). MinTEs RTs performances on the training and test sets are shown in different colours.

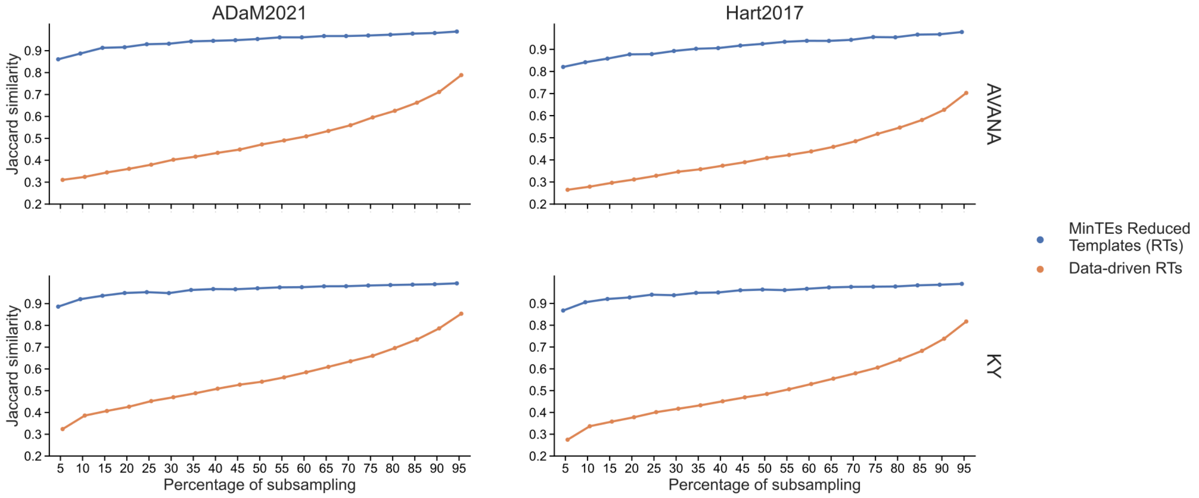


**Fig. S3 - Performances of MinTEs library-specific vs DepMap-data-driven Reduced Templates (RTs).** Average Jaccard Similarity scores, of fitness gene-sets obtained with BAGEL when using as classification templates the MinTEs derived RTs or data-driven selected RTs based on the DepMap datasets (as indicated by the different colours) obtained from two state-of-the-art reference essential gene-sets (different columns) and the same reference set of nonessential genes), for two different single guide RNA libraries (different rows), across percentage of subsampling. For DepMap-data-driven RTs, the bars show 95% confidence interval computed across 1,000 train-test reshufflings of the input dataset for each combination of parameters (i.e. reference gene-set, CRISPR-Cas9 library and percentage of subsampling).

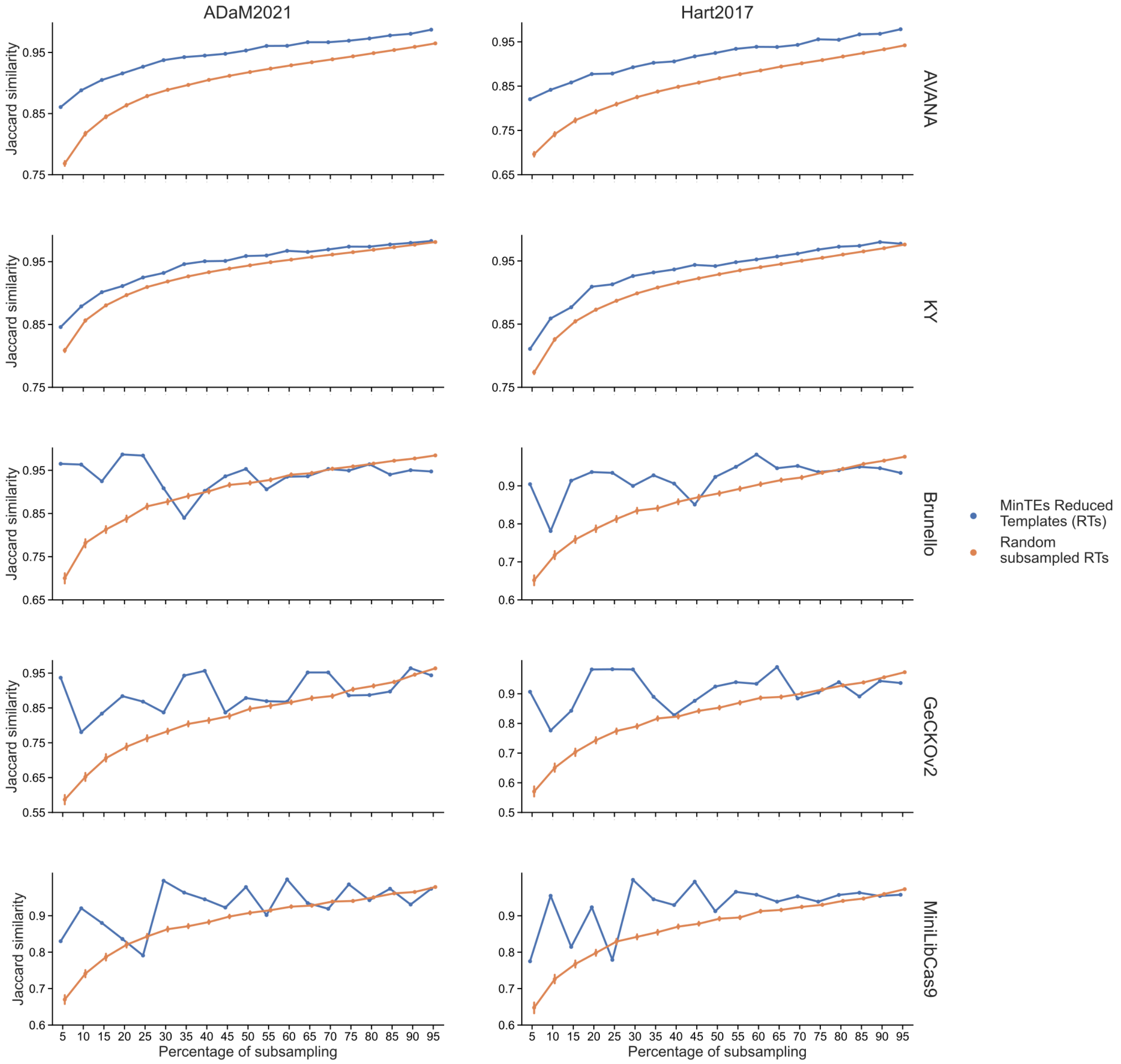


**Fig. S4 - Performances of MinTEs library-independent vs random Reduced Templates (RTs) using BAGEL.** Average Jaccard Similarity scores, of fitness gene-sets obtained with BAGEL when using as classification templates the MinTEs derived RTs or randomly selected RTs (as indicated by the different colours) obtained from two state-of-the-art reference essential gene-sets (different columns) and the same reference set of non essential genes), for five different single guide RNA libraries (different rows), across percentage of subsampling. For random performances, the bars show 95% confidence interval computed across 1,000 train-test reshufflings of the input dataset for each combination of parameters (i.e. reference gene-set, CRISPR-Cas9 library and percentage of subsampling).

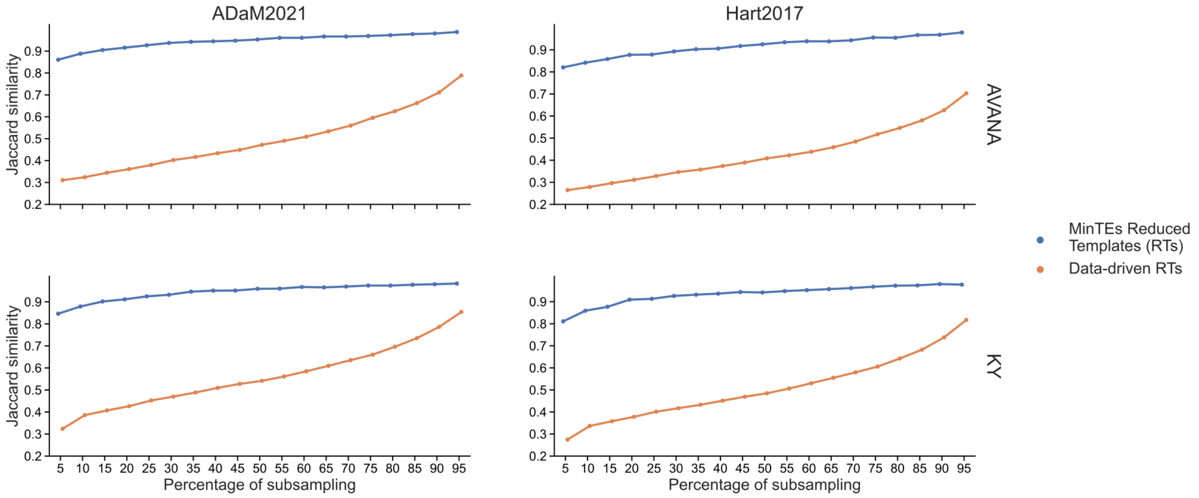


**Fig. S5 - Performances of MinTEs library-independent vs DepMap-data-driven Reduced Templates (RTs) using BAGEL.** Average Jaccard Similarity scores, of fitness gene-sets obtained with BAGEL when using as classification templates the MinTEs derived RTs or data-driven selected RTs based on the DepMap datasets (as indicated by the different colours) obtained from two state-of-the-art reference essential gene-sets (different columns) and the same reference set of nonessential genes), for two different single guide RNA libraries (different rows), across percentage of subsampling. For DepMap-data-driven RTs, the bars show 95% confidence interval computed across 1,000 train-test reshufflings of the input dataset for each combination of parameters (i.e. reference gene-set, CRISPR-Cas9 library and percentage of subsampling).

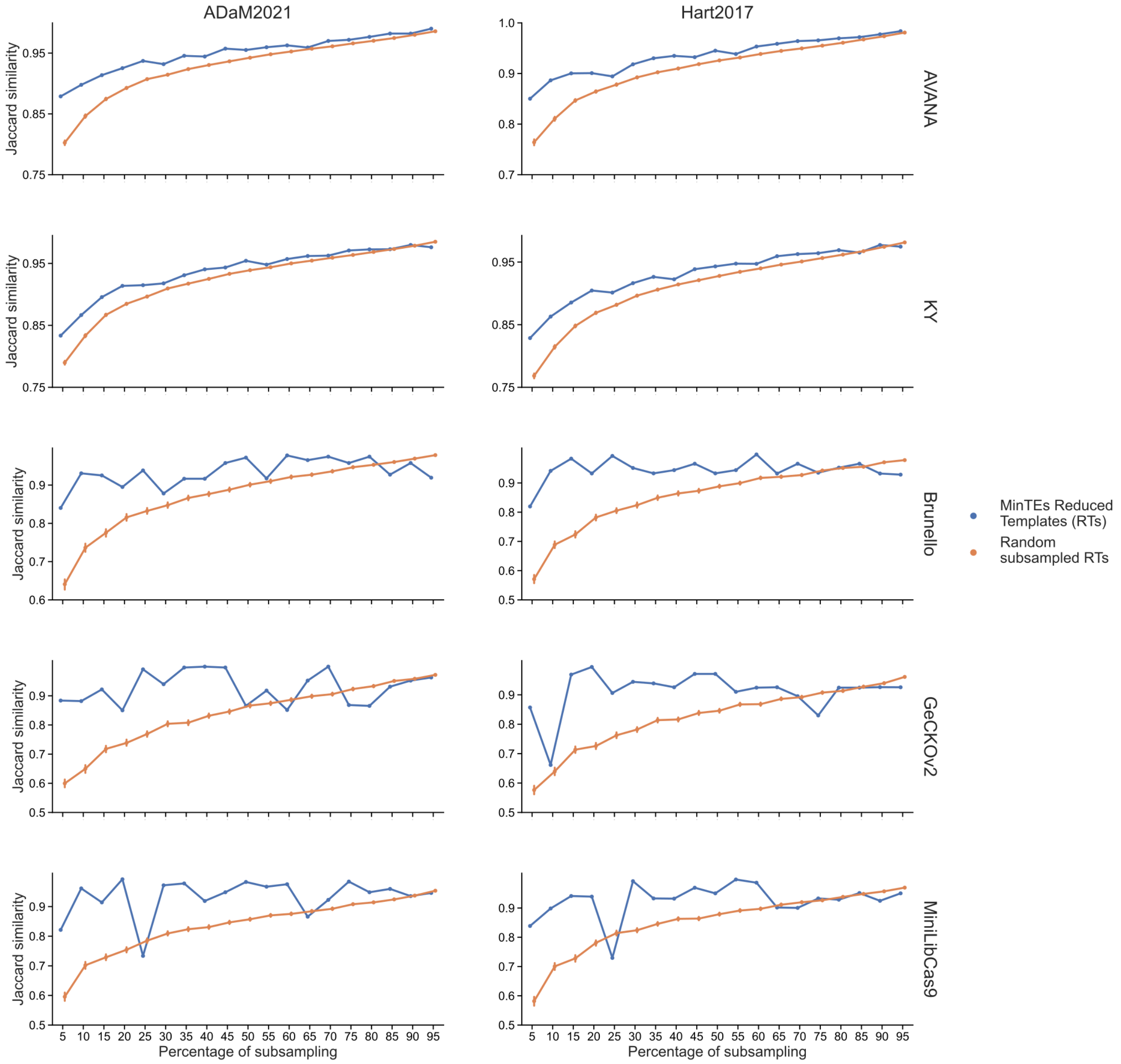


**Fig. S6 - Performances of MinTEs library-independent vs random Reduced Templates (RTs) using the False Discovery Rate (FDR) method.** Average Jaccard Similarity scores, of fitness gene-sets obtained with the FDR method when using as classification templates the MinTEs derived RTs or randomly selected RTs (as indicated by the different colours) obtained from two state-of-the-art reference essential gene-sets (different columns) and the same reference set of non essential genes), for five different single guide RNA libraries (different rows), across percentage of subsampling. For random performances, the bars show 95% confidence interval computed across 1,000 train-test reshufflings of the input dataset for each combination of parameters (i.e. reference gene-set, CRISPR-Cas9 library and percentage of subsampling)..

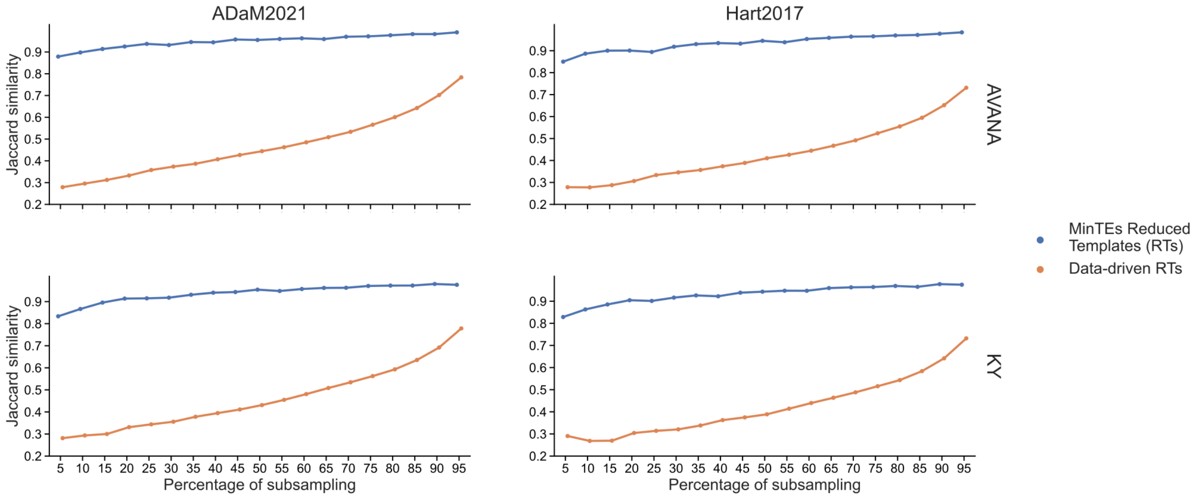


**Fig. S7 - Performances of MinTEs library-independent vs DepMap-data-driven Reduced Templates (RTs) using the FDR method.** Average Jaccard Similarity scores, of fitness gene-sets obtained with the FDR method when using as classification templates the MinTEs derived RTs or data-driven selected RTs based on the DepMap datasets (as indicated by the different colours) obtained from two state-of-the-art reference essential gene-sets (different columns) and the same reference set of nonessential genes), for two different single guide RNA libraries (different rows), across percentage of subsampling. For DepMap-data-driven RTs, the bars show 95% confidence interval computed across 1,000 train-test reshufflings of the input dataset for each combination of parameters (i.e. reference gene-set, CRISPR-Cas9 library and percentage of subsampling).

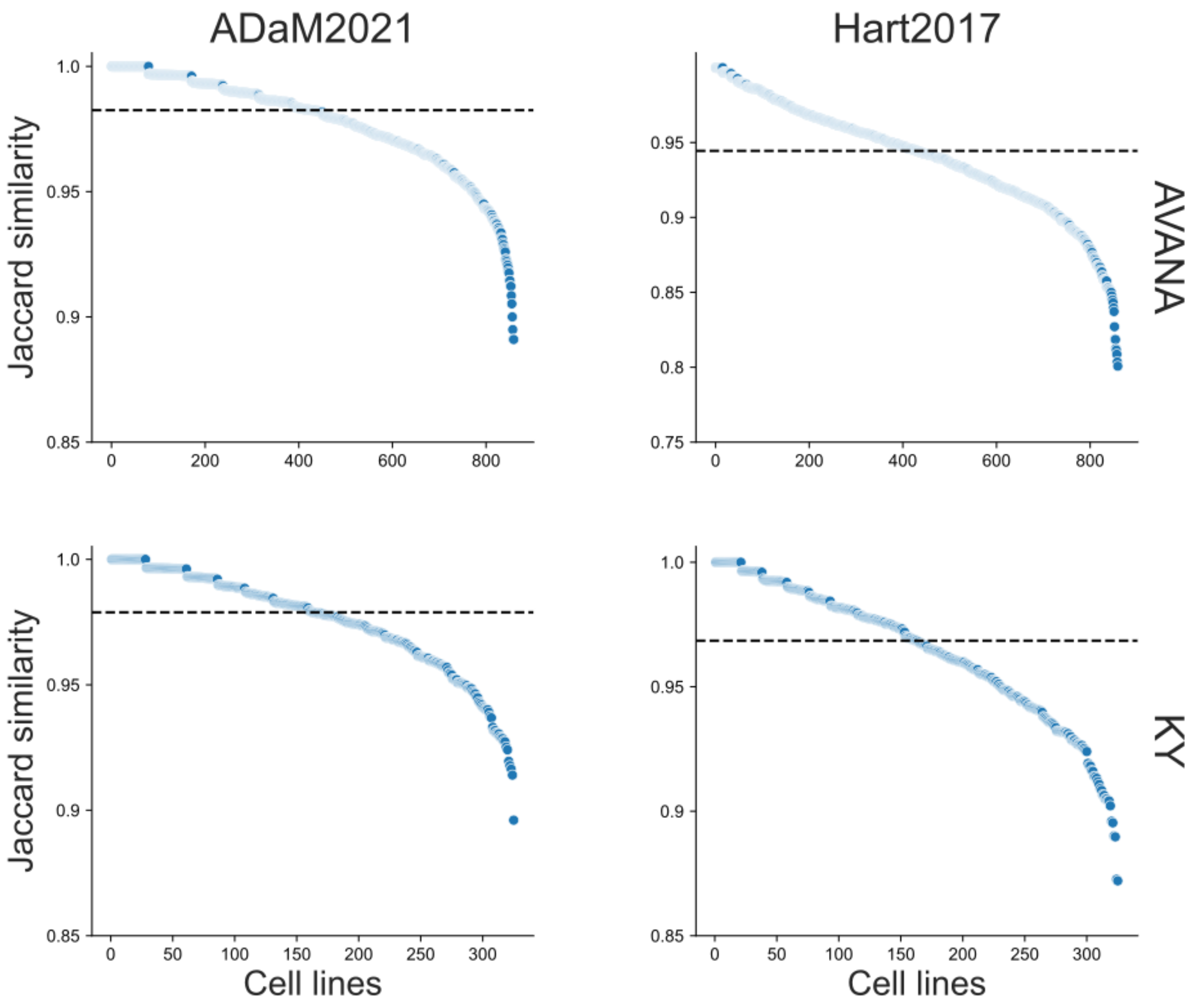


**Fig. S8 - Depletion fold-change scaling using MinTEs library-independent Reduced Templates (RTs) focused on independent essential genes**. In all the plots, points indicate individual cell line screens. Jaccard similarity scores between essential gene-sets called post scaling for two different single guide RNA libraries (different rows) with the whole reference gene-sets and the MinTEs 5% reduced RTs, respectively, with different reference essential gene-sets and the same reference set of nonessential genes (different columns). Dashed lines indicate median values. The Jaccard similarity scores are derived considering only the independent reference essential gene-sets.

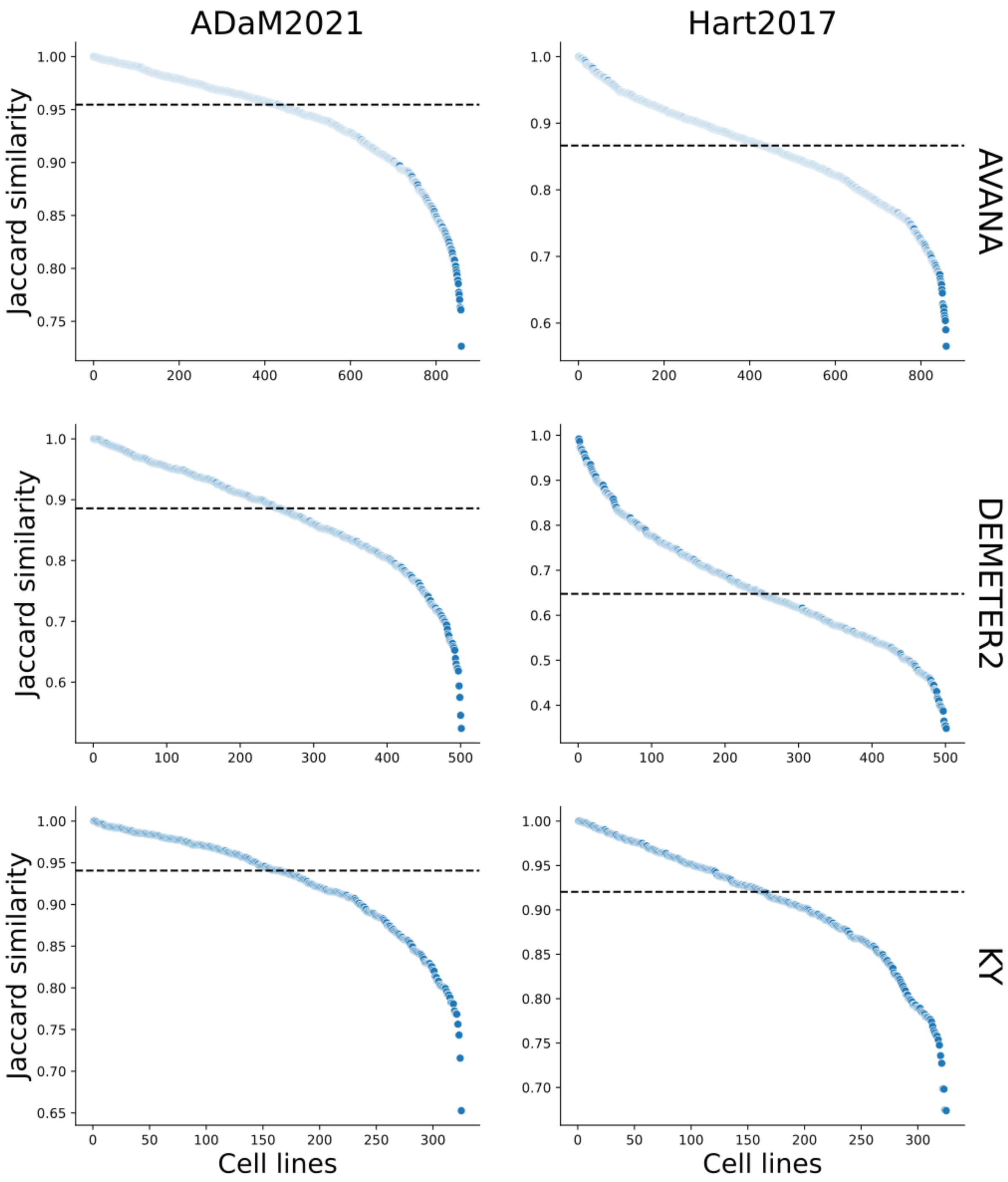


**Fig. S9 - Depletion fold-change scaling using MinTEs library-independent Reduced Templates (RTs)**. In all the plots, points indicate individual cell line screens. Jaccard similarity scores between all of the fitness genes called post scaling for DEMETER2 and two different single guide RNA libraries (different rows) with the whole reference gene-sets and the MinTEs 5% reduced RTs, respectively. Plots in different columns indicate different reference gene-sets. The Jaccard similarity scores are derived considering all the fitness genes < -0.5. Dashed lines indicate median values.

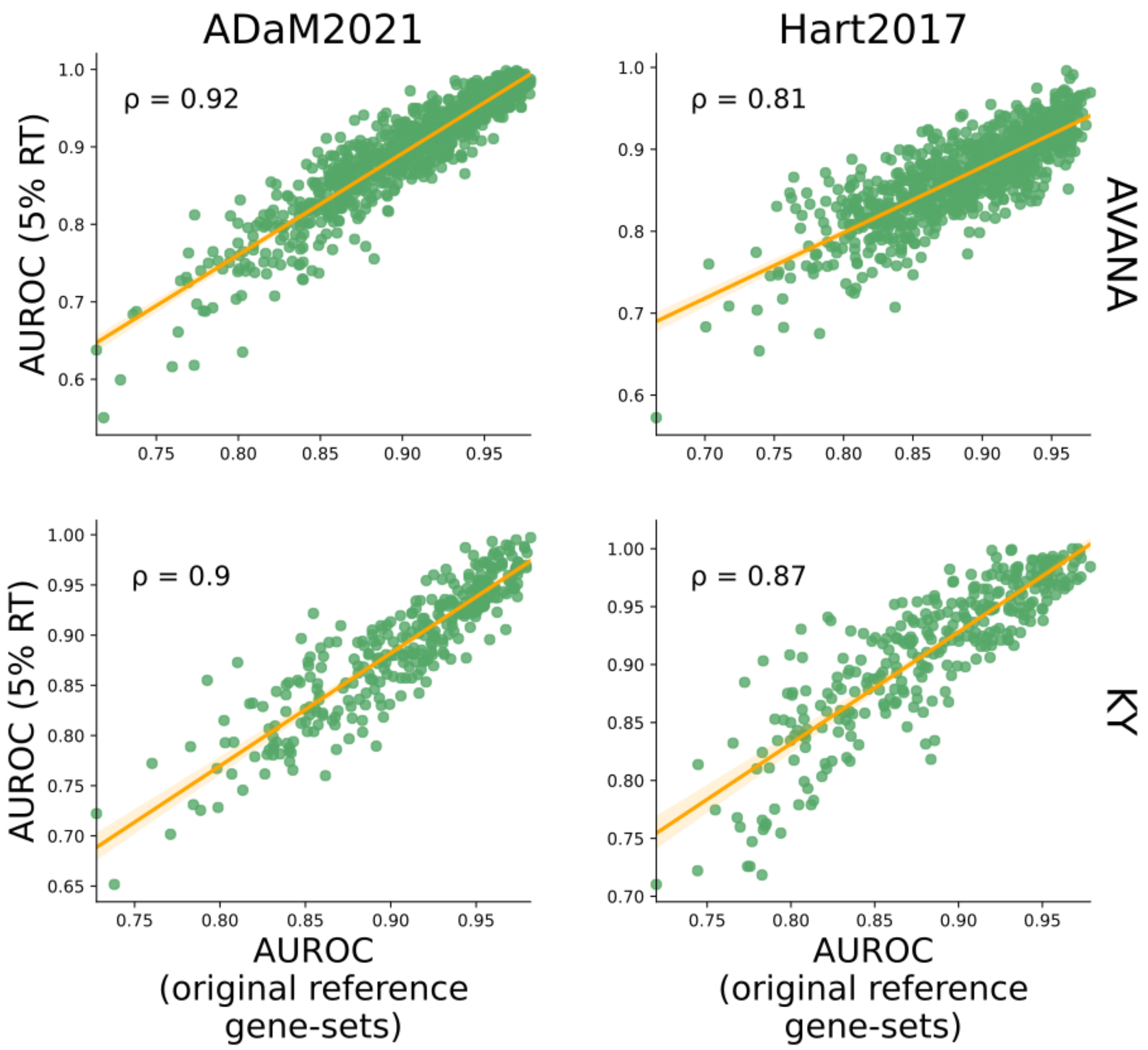


**Fig. S10 - Quality control assessment, measured in terms of Area Under the Receiver Operating Characteristics (AUROC), using MinTEs library-independent Reduced Templates (RTs)**. In all the plots, points indicate individual cell line screens. AUROC scores computed on two different single guide RNA libraries (different rows) using the whole reference gene-sets (x-axis) or the MinTEs 5% RTs (y-axis), respectively. Plots in different columns indicate different reference gene-sets. On the top left, we show the Pearson correlation coefficient. We applied a noise injection strategy to move essential and nonessential original reference gene-sets' distributions closer to each other in a randomised manner, in order to simulate low- and middle-quality screens.

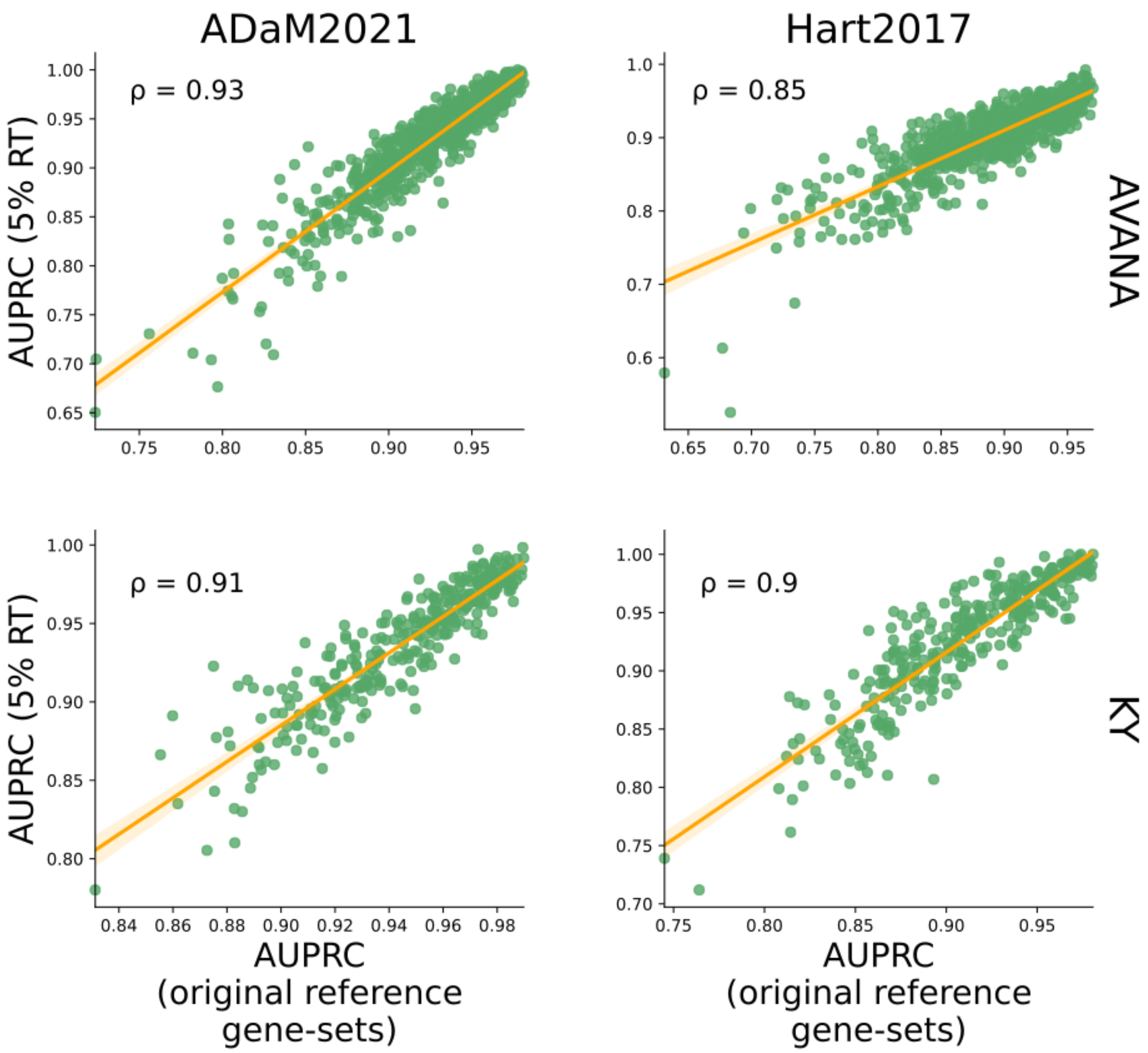


**Fig. S11 - Quality control assessment, measured in terms of Area Under the Precision Recall Curve (AUPRC), using MinTEs library-independent Reduced Templates (RTs)**. In all the plots, points indicate individual cell line screens. AUPRC scores computed on two different single guide RNA libraries (different rows) using the whole reference gene-sets (x-axis) or the MinTEs 5% RTs (y-axis), respectively. Plots in different columns indicate different reference gene-sets. On the top left, we show the Pearson correlation coefficient. We applied a noise injection strategy to move essential and nonessential original reference gene-sets' distributions closer to each other in a randomised manner, in order to simulate low- and middle-quality screens.

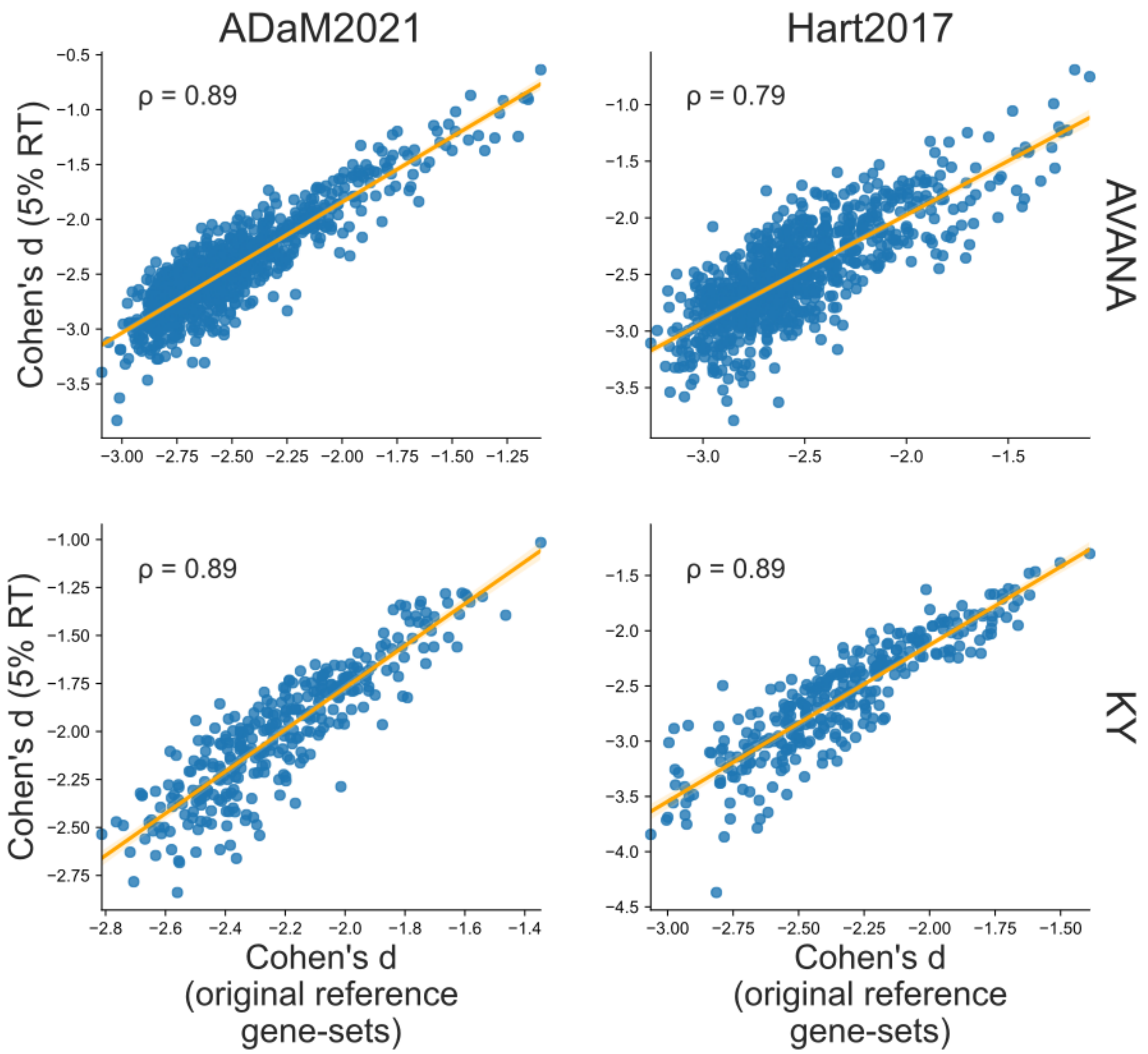


**Fig. S12 - Quality control assessment, measured in terms of Cohen's d, using MinTEs library-independent Reduced Templates (RTs)**. In all the plots, points indicate individual cell line screens. Cohen's d scores computed on two different single guide RNA libraries (different rows) using the whole reference gene-sets (x-axis) or the MinTEs 5% RTs (y-axis), respectively. Plots in different columns indicate different reference gene-sets. On the top left, we show the Pearson correlation coefficient.

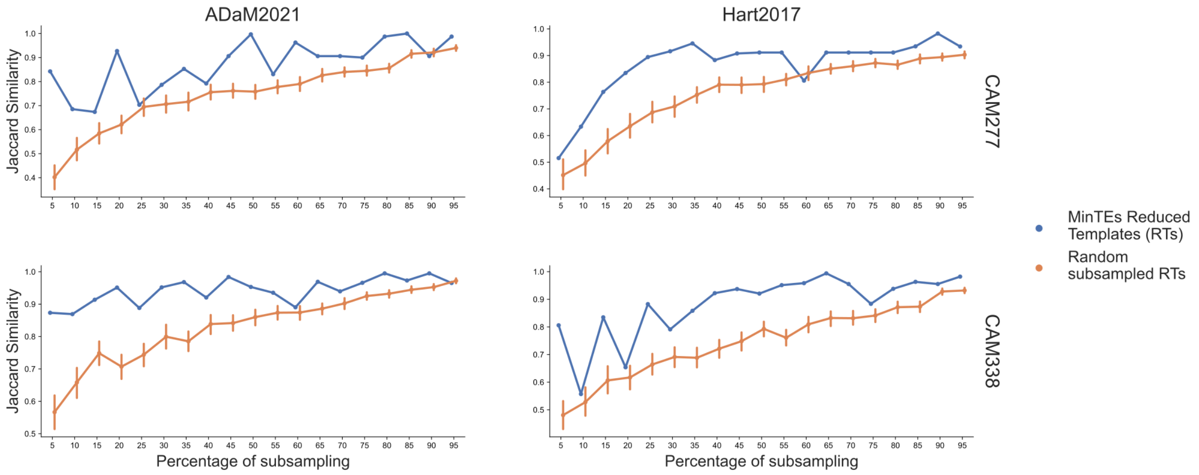


**Fig. S13 - Performances of MinTEs library-independent vs random Reduced Templates (RTs) on organoids.** Average Jaccard Similarity scores, of fitness gene-sets obtained with BAGEL when using as classification templates the MinTEs derived RTs or randomly selected RTs (as indicated by the different colours) obtained from two state-of-the-art reference essential gene-sets (different columns) and the same reference set of non essential genes), for two different organoid samples derived from oesophageal cancer (different rows), across percentage of subsampling. For random performances, the bars show 95% confidence interval computed across 1,000 train-test reshufflings of the input dataset for each combination of parameters (i.e. reference gene-set, CRISPR-Cas9 library and percentage of subsampling).
